# Arabidopsis phenotyping through Geometric Morphometrics

**DOI:** 10.1101/194449

**Authors:** Carlos A. Manacorda, Sebastian Asurmendi

## Abstract

In recent years, much technical progress has been done regarding plant phenotyping including the model species *Arabidopsis thaliana*. With automated, high-throughput platforms and the development of improved algorithms for the rosette segmentation task, it is now possible to massively extract reliable shape and size parameters for genetic, physiological and environmental studies. The development of low-cost phenotyping platforms and freeware resources make it possible to widely expand phenotypic analysis tools for Arabidopsis. However, objective descriptors of shape parameters that could be used independently of platform and segmentation software used are still lacking and shape descriptions still rely on *ad hoc* or even sometimes contradictory descriptors, which could make comparisons difficult and perhaps inaccurate. Modern geometric morphometrics is a family of methods in quantitative biology proposed to be the main source of data and analytical tools in the emerging field of phenomics studies. It has been used for taxonomists and paleontologists for decades and is now a mature discipline. By combining geometry, multivariate analysis and powerful statistical techniques, it offers the possibility to reproducibly and accurately account for shape variations amongst groups. Based on the location of homologous landmarks points over photographed or scanned specimens, these tools could identify the existence and degree of shape variation and measure them in standard units. Here, it is proposed a particular scheme of landmarks placement on Arabidopsis rosette images to study shape variation in the case study of viral infection processes. Several freeware-based geometric morphometric tools are applied in order to exemplify the usefulness of this approach to the study of phenotypes in this model plant. These methods are concisely presented and explained. Shape differences between controls and infected plants are quantified throughout the infectious process and visualized with the appealing graphs that are a hallmark of these techniques and render complex mathematical analysis simple outcomes to interpret. Quantitative comparisons between two unrelated ssRNA+ viruses are shown and reproducibility issues are assessed. Combined with the newest automatons and plant segmentation procedures, geometric morphometric tools could boost phenotypic features extraction and processing in an objective, reproducible manner.

## Introduction

Plant phenotyping is the process of recording quantitative and qualitative plant traits. It is key to study plant responses to the environment (Granier and Vile, 2014).

A 2016 IPPN survey (https://www.plant-phenotyping.org/ippn-survey_2016) between plant scientists found that most participants think that plant phenotyping will play an important role in the future, being stress assessing and the model plant *Arabidopsis thaliana* mentioned between the topics of main interest.

Recently, many new techniques have been developed to facilitate and improve quantitative plant phenomics (i.e. the full set of phenotypic features of an individual), going from destructive to non-destructive and even high-throughput phenotyping (Vanhaeren et al., 2015). Whereas the throughput is an important aspect of phenotyping, spatial and temporal resolutions, as well as accuracy, should be considered (Dhondt et al., 2013).

Several workers (Camargo et al., 2014; De Vylder et al., 2012; Green et al., 2012; Ispiryan et al., 2013; Tessmer et al., 2013) have developed freely available software that overcome the difficult task of rosette segmentation (an issue still under investigation) by different means. This software allows several rosette parameters to be computed such as area and perimeter in addition to other more complex descriptors. However, the persistence of *ad hoc* descriptors (Bonhomme et al., 2013; Krieger, 2010) and the lack of a gold standard in this actively developing field, could give rise to reproducibility issues, due to different growing substrate-segmentation algorithms combinations. Moreover, different approaches give sometimes the same name to different parameters (e.g. “roundness” in ImageJ (Schneider et al., 2012) vs. (Camargo et al., 2014)) or different names to the same parameter (e.g. “solidity” in (Ispiryan et al., 2013) equals “compactness” in (Camargo et al., 2014; De Vylder et al., 2012) and “surface coverage” in (Vanhaeren et al., 2015)). The need of developing objective, mathematically and statistically sound and more accurate shape descriptors in plants has been stressed out in recent reviews on the topic (Balduzzi et al., 2017; Bucksch et al., 2017; Lobet, 2017).

Nonetheless, image datasets analyses require a conceptual and statistical corpus of knowledge that is not always present in a plant biologist’s research field. Plant phenotyping relies on skills and technologies that are used to characterize qualitative or quantitative traits regardless of the throughput of the analyses (Granier and Vile, 2014). One such knowledge corpus is morphometrics (Strauss, 2010).

Traditional morphometric analyses such as measures and ratios of length, depth and width were widely used in Paleontological and Zoological studies throughout the 20^th^ century. To the end of that century the seminal work of (Thompson, 1917) was re-evaluated under the light of multivariate analysis and novel mathematical developments (Kendall, 1977; Kendall and Kendall, 1980), giving rise to modern geometric morphometrics (GM), in which was called a “revolution” in morphometrics (Adams et al., 2004; Bookstein, 1996b; James Rohlf and Marcus, 1993).

GM combines geometry, multivariate morphometrics, computer science and imaging techniques for a powerful and accurate study of organismal forms. This family of methods in quantitative biology is proposed to be the main source of data and analytical tools in the emerging field of phenomics (Cardini and Loy, 2013). Formally, GM is “a collection of approaches for the multivariate statistical analysis of Cartesian coordinate data, usually (but not always) limited to landmark point locations” (http://life.bio.sunysb.edu/morph/glossary/gloss1.html). Landmark methods have been successfully applied to various species, and have the advantage of being easy to understand (Cope et al., 2012).

Besides enhanced statistical power and better descriptive and graphical tools, GM allow researchers to decompose form in size and shape, and the whole configuration of the organism under study is analyzed, rather relying on the description of relative displacements of pairs of points.

GM is now a mature discipline that has been widely applied in biology (Claude, 2013; MacLeod et al., 2013; Zelditch et al., 2000) (see (Cardini, Rohlf, Klingenberg, Adams, 2013) for a review). In plants, leaves of grapevine (Chitwood et al., 2016) and oak (Viscosi, 2015; Viscosi and Cardini, 2011) were studied using GM methods.

Plant viruses cause important worldwide economic losses in crops (Scholthof et al., 2011). Symptoms include plant stunting, changes in leaf morphology, and sometimes plant death (Matthews et al., 2002) and vary depending on various aspects including genetic compatibility and environmental conditions.

Even given a particular host-virus interaction, different viral strains trigger different symptomatology, which are more or less subtle for the observer to distinguish (Manacorda et al., 2013; Sánchez et al., 2015; Zavallo et al., 2015). Comparing the severity of qualitative viral symptoms is a difficult task performed mainly by visually rating symptoms (e.g. (Doumayrou et al., 2013)). Consequently, morphological differences could be difficult to describe and reproducibility issues could arise.

*Arabidopsis thaliana* (L.) Heynh. has been extensively used in studies of influences of environmental factors on plants, paving the path to the development and testing of experimental techniques or data analysis methods (Ferrier et al., 2011). The Arabidopsis rosette is a nearly two-dimensional structure in the vegetative phase (Ispiryan et al., 2013), which facilitates image acquisition and interpretation.

Here, it is proposed a case study where GM tools are applied to study and quantitatively describe morphometric changes triggered in *A. thaliana* plants by RNA viruses belonging to two unrelated families. It is proposed a particular selection of landmarks to locate in the Arabidopsis rosette during its vegetative phase. The study spans from the earlier stages of viral infection to later ones, when symptoms are already visually detectable by naked eye. Comparisons are made between discriminant power of computer-assisted classification and expert human eye. Symptoms severity provoked by both viruses is also compared, based on the relative morphometric changes induced relative to healthy controls. Changes in allometric growth, phenotypic trajectories and morphospace occupation patterns are also investigated. Size analyses are also performed and the problem of growth and development modeling is discussed in the context of viral infections. Throughout this work, several bioinformatics resources are applied, in order both to extract the higher degree of information available, but also to exemplify different and complementary possibilities that nowadays GM offers for the accurate description of shape in Arabidopsis.

## Materials and Methods

### Plant growth conditions

*A. thaliana* Col-0 seeds were stratified at 4°C for 3 days. Plants were grown under short days conditions (10 h light/14 h dark cycle, T(°C)= 23/21, Hr(%)= 60/65, and a light intensity of 150 μE m-2 s-1) in a controlled environmental chamber (Conviron PGR14; Conviron, Winnipeg, Manitoba, Canada). Plants were grown in individual pots in trays and treatments were assigned to plants in all trays. One experiment was performed with ORMV and two independent experiments were carried on with TuMV-UK1.

### Virus infection assays

ORMV (Oilseed Rape Mosaic Virus) (Aguilar et al., 1996) was maintained in Nicotiana tabacum (cv. Xhanti nn) and infective sap was obtained after grinding infected leaves with mortar and pestle in 50 mM phosphate buffer pH=7.5. TuMV (Turnip Mosaic Virus)-UK1 strain (accession number X65978) (Sánchez et al., 1998) was maintained in infected A. thaliana Col-0. Fresh sap was obtained immediately prior to use to inoculate plants with sodium sulfite buffer (1% K2HPO4 + 0,1% Na2SO3 [wt/vol]). Mock-inoculated plants were rubbed with carborundum dust with either 50 mM phosphate buffer pH=7.5 or sodium sulfite buffer, respectively. Plants were mechanically inoculated in their third true leaf at stage 1.08 at 21 DPS, (Boyes et al., 2001) because those leaves were almost fully developed by the time of the procedure and therefore constituted a source tissue for the export of virions to the rest of the plant.

### Image acquisition

Zenithal photographs of individual plants growing in pots were taken with a Canon PowerShot SX50HS camera mounted in a monopod at maximal resolution. Photographs were taken at the same time of the day in successive days to minimize error. A ruler was placed in each image acquisition and only its central part (60-80 mm) was taken into account to avoid image distortion at the edges of the photograph (Schutz and Krieger, 2007).

### Landmark configuration and digitization

Specimens were imaged at each DPI and .JPG files were opened with TPSUtil software, a member of the TPS Series of GM tools (Rohlf, 2015, 2017) that prepares the data for further analyses. Opening the output .TPS files with with TPSDig2 is the first step to digitization of landmarks. The 11 landmarks were digitized in the same order on each picture, after setting a scale factor with a ruler, at each DPI. Eleven landmarks were recorded for each plant. Landmarks were selected to fulfill the basic requirements for 2D approximation (Zelditch et al., 2012) (see main text). Following (Bookstein, 1991) criteria, landmark 11 (which is situated at the centre of the rosette) is a Type 1 landmark. Landmarks 1, 2, 3, 4 and 5 (which are located at the tip of leaves #8 to #12 and are the maxima of curvature of that structure) and landmarks 6, 7, 8, 9 and 10 (which are located at the intersection of the petiole and the lamina of each leaf from #8 to #12) cannot be unambiguously assigned due to the continuous nature of the leaf curvature and are Type 2 landmarks. Each specimen was digitized in less than 1 minute. The Output of TPSDig2 is a .TPS file containing information about specimen name, scale factor, and raw coordinates of each landmark for all specimens digitized. Landmark digitization was repeated to estimate measurement error for each specimen a week after the first digitization.

### Validation of the tangent space approximation

For a given M-dimensional structure with K landmarks (here, M = 2 and K = 11) it can be imagined an individual´s shape as a point in an M X K multidimensional space (a hypersphere). After centering and rescaling, 3 dimensions are lost and shapes are said to be in a pre-shape space; they are not rotated yet. The distance in the surface of the hypersphere at which rotation differences between shapes are minimal, is called Procrustes distance. Afterwards, a reference (average) shape is selected and all other shapes are rotated to minimize distances relative to it, generating a shape space and losing one more dimension (remaining 2K-4). Because distances over curved multidimensional spaces are non-Euclidean, conventional tools of statistical inference cannot be used. Fortunately, for most biological shapes an approximation to Euclidean distances is valid, by projecting shape points to a tangent Euclidean space (for a visual explanation see (Zelditch et al., 2012)). This assumption should, however, be tested when new forms are being analyzed. The TPSSmall program is used to determine whether the amount of variation in shape in a data set is small enough to permit statistical analyses to be performed in the linear tangent space approximate to Kendall's shape space which is non-linear. Since TPSSmall does not perform reflections, datasets analyzed with TPSDig2 were opened again and specimens reflected when necessary to leave all clockwise rosettes.

### Statistical analyses

Except otherwise stated, shape analyses were performed using MorphoJ (Klingenberg, 2011) and the TPS series (Rohlf, 2015), as described in the main text.

Student´s t, Mann-Whitney, paired Hotelling´s tests and rosette growth parameters analyses were executed in PAST (Hammer et al., 2001).

PTAs based on (Adams and Collyer, 2009) were run in R (RStudio Team, 2016).

Linear and nonlinear regressions for growth and development modeling and analysis of residuals autocorrelations were performed in PAST and Infostat (Di Rienzo et al., 2012), since both software yield complementary information. Lack-of-fit tests were performed in Infostat.

Excel 2010 was used for Durbin-Watson panel test, Holm´s-Bonferroni sequential test for multiple comparisons (Gaetano, 2013; Holm, 1979) and hyperellipses calculations using Real Statistics for Excel 2010 (ver. 4.14) (Zaiontz, 2017).

## Results

This work aims to introduce the use of GM tools for the analysis of Arabidopsis rosette phenotypes in an objective and repeatable way. As such, it is not intended to offer a complete introductory explanation of each GM tool, an objective that is beyond the scope of this paper. Such a task was already performed by (Viscosi and Cardini, 2011) and for a complete introductory explanation of GM tools applied in biological systems it is recommended the lecture of (Zelditch et al., 2012). Software used in this work frequently has its own user´s manual and informative examples (Di Rienzo et al., 2012; Hammer et al., 2001; Klingenberg, 2011; Rohlf, 2015). Nevertheless, with the purpose to facilitate the comprehension of this work to newcomers in the field of GM, each tool is briefly described prior to its application throughout the Results section.

Morphometrics aims at analyzing the variation and covariation of the size and shape of objects, defining altogether their form. Shape and form might be confusing words, used as synonyms in many languages (Bonhomme et al., 2013). Hereafter, it will be followed the GM definition of shape in the sense of (Kendall, 1977) that it is “all the geometric information that remains when location, scale and rotational effects are filtered out from an object”.

### Landmarks digitization, Procrustes fit and outliers detection

At the heart of GM analyses is the concept of landmarks. Landmarks are discrete anatomical *loci* that can be recognized as the same point in all specimens in the study. They are homologous points both in an anatomical and mathematical sense. The selection of landmarks is based in the observance of five basic principles (Zelditch et al., 2012):

1) Homology. Landmarks are sequentially numerated and each landmark must correspond to the same number of landmark in all specimens under study.
2) Adequate Coverage of the Form. Landmarks should be chosen in a way they cover up the maximum possible extent of the form of interest. It is important to bear in mind that a region not included between landmarks is not analyzed.
3) Repeatability. The same landmarks should be easily identified in the same structure in order to avoid measurement error.
4) Consistency of Relative Position. This attribute guarantees that landmarks do not interchange relative positions.
5) Coplanarity. For 2D-landmarks, an assumption of coplanarity is required to avoid measurement error.

There is no absolute landmark configuration on any given structure. The choice of the number of landmarks and their configuration depend on the hypotheses being tested. Here, the focus of the analyses is on the phenotypic impact of viral infections on the Arabidopsis rosette through time. Hence, short-day conditions were chosen to delay flowering, allowing the plant´s aerial part to remain near two-dimensional during the experiment. To encompass as broadly as possible the phenotypic changes experienced by the plant during the infection, chosen landmarks should not only be present from earlier stages to the infection to later ones, but also be placed in regions that experience dramatic changes upon infection. A relatively reduced number of landmarks can be used to describe complex forms (Carreira et al., 2011; Viscosi and Cardini, 2011).

An 11-landmark configuration for the Arabidopsis rosette is shown in Figure 1A (see Materials and Methods). Plants were inoculated in their third true leaf (24 plants were mock-inoculated and 17 were ORMV-infected) and images were acquired starting from three days post-inoculation (DPI) to 12 DPI (see Materials and Methods). Leaves below number 8 were not chosen for landmark placement for three main reasons: a) they are hidden for younger leaves at later stages of infection b) these old leaves had almost finished their growth by the time the first photographs were taken (and the form covered would be a less informative one for the process of shape and size change upon viral infection) and c) the senescence process of older leaves lead to morphological changes derived from dehydration and death. Younger leaves (beyond leaf number 12) were not chosen because they were not present at the earlier stages of infections, therefore violating the requisite of repeatability of landmarks.

**Figure 1.**
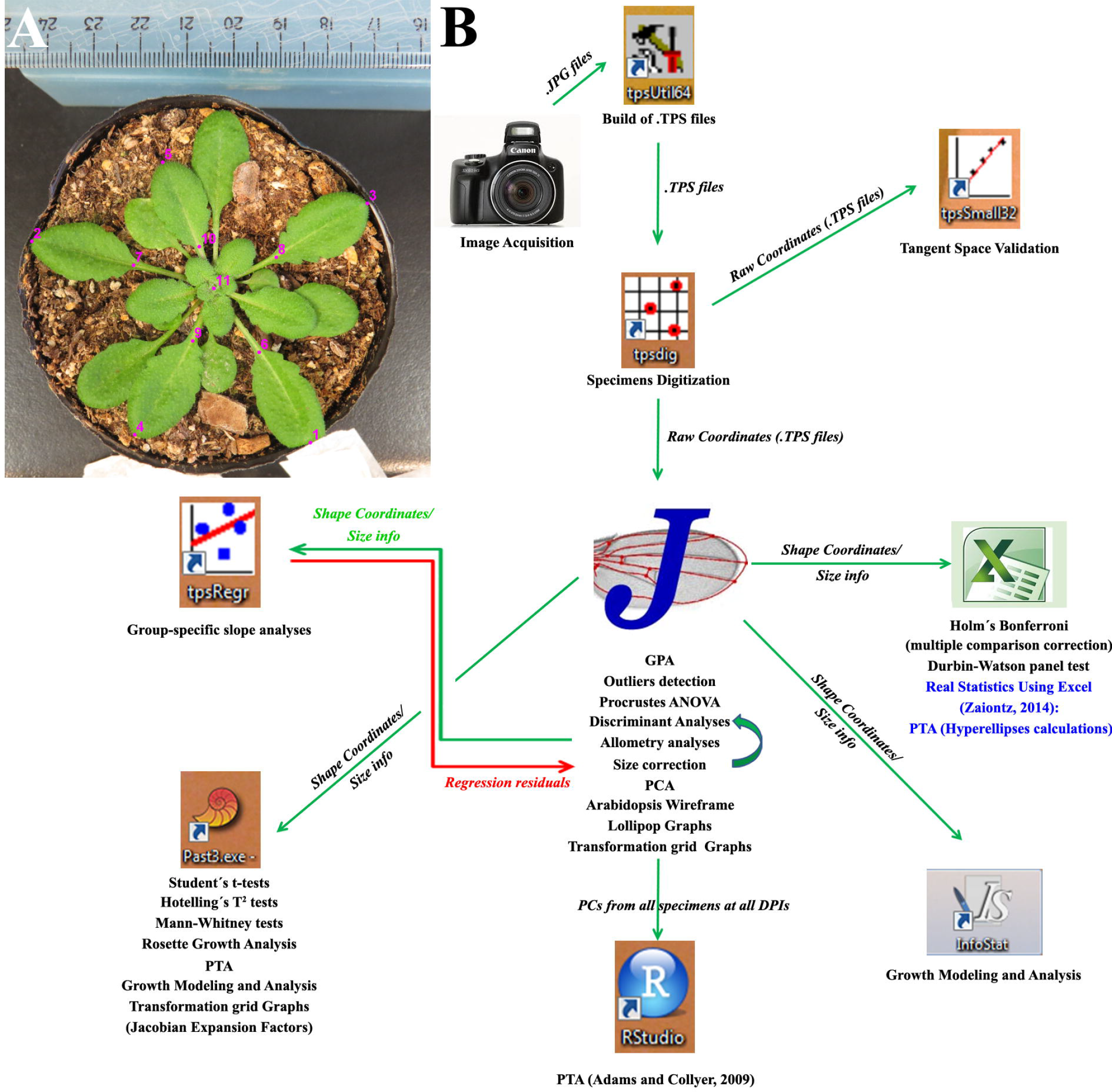
**(A)** Landmark configuration in an Arabidopsis rosette. An 8 DPI Mock-inoculated rosette is shown. (B) Analysis flowchart showing the different software used in this study, with main features extracted from each one listed below corresponding icon. See main text and Materials and Methods for details.

A flowchart of data analyses throughout this paper is shown in Figure 1B. Image datasets for all DPIs and both treatments were handled and digitized for further analyses using the TPSUtil and TPSDig2 software generating .TPS output files. Digitization process was performed twice (see Materials and Methods).

Several freeware could be used to extract shape information from .TPS files (Zelditch et al., 2012). Here, the MorphoJ software (Klingenberg, 2011) was chosen mainly because of its ease to use and comprehensive tools available. MorphoJ creates new datasets from several file extensions, including .TPS. The “Supplementary file ORMV.morphoj” was created and 16 datasets were generated, one for each DPI and digitization instance. Specimens were classified according with ID, Treatment, DPI and Digitization for each dataset. Combinations of classifiers were also done to perform further grouped analyses.

The first step of shape analysis in GM consists in extracting shape coordinates from raw data obtained at the digitization step. The procedure that has become standard in GM studies is the Generalized Procrustes Analysis (GPA).

The purpose of Procrustes procedures is to remove from the specimens all information that is not relevant for shape comparisons, including size. Specimens are firstly translated at the origin (“superimposed”) by subtracting the coordinates of its centroid from the corresponding (X or Y) coordinates of each landmark. Then, differences in size are removed rescaling each specimen to the mean centroid size (CS) (CS is calculated as the square root of the summed squared distances of each landmark from the centroid, giving a linearized measure of size). Differences in rotation are eliminated by rotating specimens minimizing the summed squared distances between homologous landmarks (over all landmarks) between the forms. The process starts with the first specimen, and then an average shape is found that now serves as a reference. It proceeds iteratively over all specimens until no further minimization of average distances are found (Rohlf and Slice, 1990). MorphoJ performs a full Procrustes fit, which is a variant of the analysis that is more conservative and resistant to outliers of shape.

In Arabidopsis, the arrangement of organs along the stem (phyllotaxy) follows a predictable pattern, the Fibonacci series. Phyllotaxy orientation can be either clockwise or counter-clockwise (Peaucelle et al., 2007). This should be taken into account because clockwise and counter-clockwise rosettes are biological enantiomorphs like right and left hands and must not be superimposed by GPA. Opportunely, MorphoJ automatically performs reflections on every specimen when executing a GPA and therefore it is not a problem at this stage, but care must be taken when using different software.

After executing a full Procrustes fit of each dataset, they were inspected for the presence of outliers. The shape of one Mock-inoculated plant (M2) diverted the most from the rest in 11 out of 16 datasets. Therefore, it was excluded from all datasets for successive analyses.

Afterwards, datasets were combined and the “Combined dataset 3-12 DPI” was created with 640 observations included following a common GPA. Then, a wireframe was created that connects consecutive landmarks. This tool aids visualization, as will be explained later. Next, the “Combined dataset 3-12 DPI” was subdivided by DPI. This creates one dataset for each DPI, each one with the two digitization outputs for each plant.

### Validation of the tangent (Euclidean) space approximation

Prior to conducting further analyses, a basic assumption of GPA-based GM analysis should be tested: that the projections of shapes in Kendall´s shape space onto a tangent Euclidean shape space are good approximations for the studied shapes. This task is performed by basically comparing the Procrustes distances (the conventional measure of a morphometric distance in geometric morphometrics (Bookstein, 1996a)) obtained using both shape spaces (see Materials and Methods). Two subsets of data were created for each DPI, one with Mock-inoculated and the other with ORMV-infected plants. Next, datasets were manually combined using a text editor to create three main datasets (Mock, ORMV and ALL plants). TPSSmall (v.1.33) was then used to compare statistics for distance to reference shape both in Tangent (Euclidean) and Procrustes (Kendall´s) shape space for both treatments separately and for all plants together (Supplementary Table 1). Results showed that maximum Procrustes distances from mean (reference) shape were 0.371 (ORMV), 0.405 (Mock) and 0.400 (ALL). They are all well below the largest possible Procrustes distance (π/2 = 1.571). Mean Procrustes distances from mean (reference) shape were 0.168 (ORMV), 0.186 (Mock) and 0.184 (ALL). This indicates a closer arrangement of ORMV shapes in shape space relative to Mock-inoculated plants. Tangent and Procrustes distances were very similar (Supplementary Table 1) and regressions through the origin for distance in tangent space, Y, regressed onto Procrustes distance, X, showed slopes > 0.98 and correlations > 0.9999 for all groups (Supplementary Table 1 and Supplementary Figure 1). This results are in line with several similar analysis performed onto a variety of biological forms (Bai et al., 2011; Dewulf et al., 2014; Parés-Casanova and Allés, 2015; Viscosi and Cardini, 2011).

### Testing measurement error and variation between treatments using Procrustes ANOVA

As mentioned before, two digitizations were performed on each plant at each DPI, in order to evaluate measurement error. This procedure is important because digitization error should always account for far less variance in the subsequent analyses than specimens and treatments do (Viscosi and Cardini, 2011). There are the differences between specimens and particularly between treatments that are worth investigating, not human error in landmark placement. Purposely, datasets for each DPI were combined and subjected to a hierarchical analysis of variance (ANOVA). In MorphoJ this is a Procrustes ANOVA, with “Treatment” as an additional main effect, “ID” for the individuals and “Digitization” as the Error1 source. In Procrustes ANOVA, variance is partitioned by means of hierarchical sum of squares (SS) which implies that each effect is adjusted for effects that appear earlier in the hierarchy. This is taking into account the nested structure of the data (an issue that is crucial if the design is unbalanced, i.e., with unequal sample sizes as is here the case), thus allowing one to quantify differences in Treatments and plants regardless of Treatment. The variance unexplained by any of these effects is measurement error and it is estimated using the differences between digitizations. Hence, total variance was decomposed into main (treatment) and random (ID and digitization) components and was expressed as a percentage of total variance for each DPI. The analysis is executed simultaneously for both size and shape. Results are shown in Supplementary Table 2. Explained variance (as a % SS) for which measurement error accounted for was in the range of 0.01 and 0.12 for size and 0.40 and 1.15 for shape over all DPIs. Thus, measurement error was negligible throughout the digitization process. Detailed analysis of results shown in Supplementary Table 2 revealed that for size, the Individual (ID) effect was highly significant at each DPI as evidenced by Goodall´s F-test (p < 0.0001). Treatment effect was insignificant from 3 to 5 DPI but starting from 6 DPI the virus affected the plant´s size (0.0001 < p < 0.03).

For shape, the Individual effect was also highly significant at each DPI as evidenced both by Goodall´s F-test (p < 0.0001) and by MANOVA (standing for Multivariate Analysis of Variance) results (p < 0.0001). Treatment impacted earlier shape than size, since as soon as 5 DPI differences in shape were detected (p < 0.001).

### Ordination Methods and shape change visualization

#### PCA

Once shape variables (the 22 Procrustes Coordinates) are extracted for all specimens at each DPI, it is useful to plot differences between individuals and treatments. However, patterns of variation and covariation between lots of variables are difficult to envision particularly because geometric shape variables are neither biologically nor statistically independent (Zelditch et al., 2012). PCA is a technique that allows simplifying those patterns and making them easier to interpret. By performing a PCA, shape variables are replaced with complex variables (principal components, PCs) that do not covary but carry all the information. Moreover, as PC axis are orthogonal and independent, and most of the variation in the sample usually can be described with only a few PCs, shape analysis could be restricted to very few axes, avoiding the need of jointly interpret dozens of variables. It is important to keep in mind that PCA is useful for the comparison between individuals, not groups, and though a powerful descriptive tool, it does not involve any statistical test. Therefore, the relative separation of groups in a PCA plot does not allow one to extract conclusions about significant differences (or its absence).

Firstly, this technique was used to inspection error measurement (previously quantified by Procrustes ANOVA, Supplementary Table 2). A covariance matrix was created for the “Combined dataset 3-12 DPI” and then a PCA was performed. Scatterplots were generated for the first 4 PCs, which together account for 87.2 % of the total variance (Figure 2). The proximity of equally colored points indicates a small digitization error.

**Figure 2.**
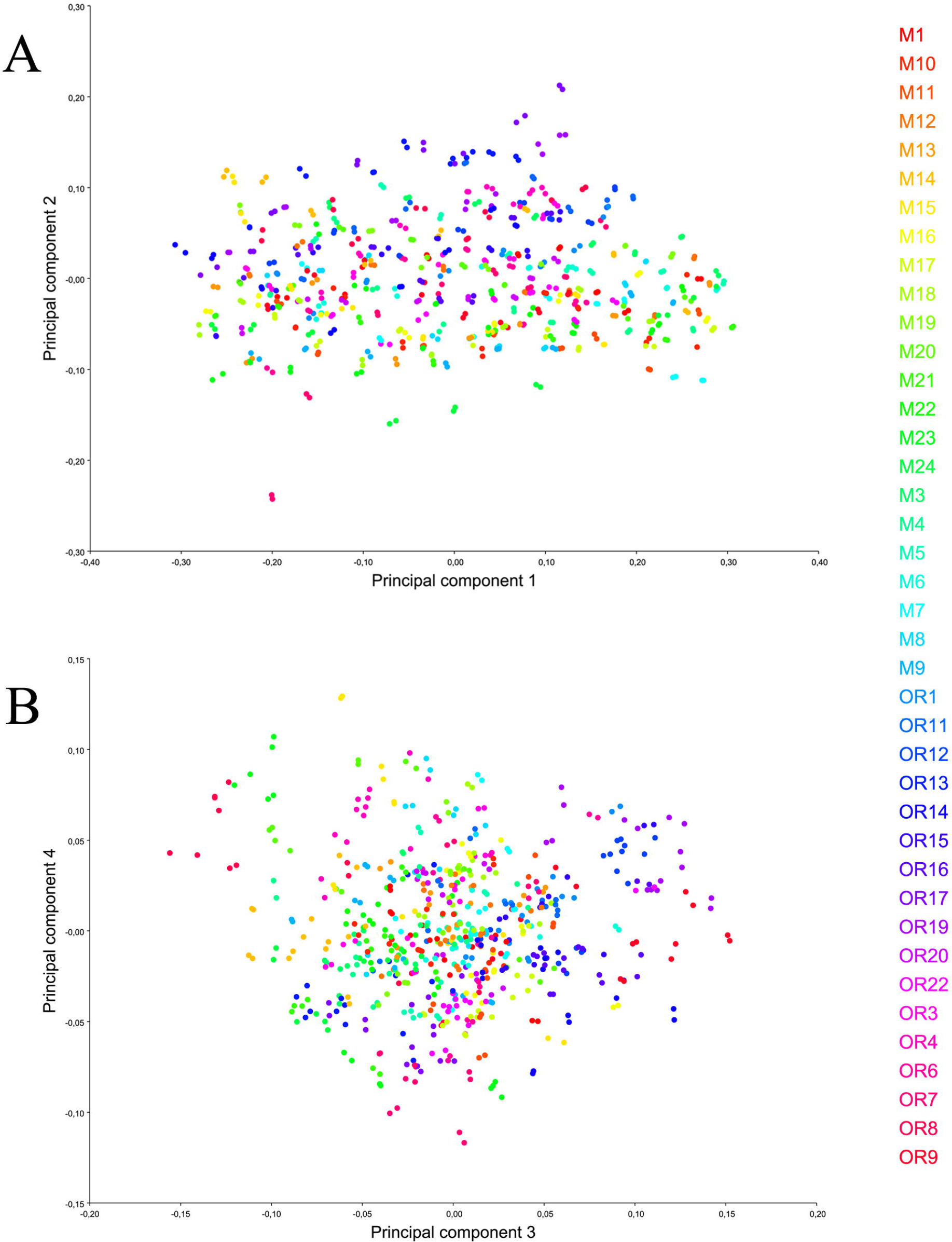
Shape variation including all observations and replicas. PCA scatterplots of (A) PC1 vs. PC2 and (B) PC3 vs. PC4. Equally colored dots represent both digitizations of the same specimen, for all DPIs. The scale factor for this graph is directly the magnitude of the shape change as a Procrustes distance; the default is 0.1, which corresponds to a change of the PC score by 0.1 units in the positive direction.

As measurement error explained a negligible percentage of variance, digitizations were averaged within specimens and DPIs. From the “Combined dataset 3-12 DPI” it was created the “Combined dataset 3-12 DPI, averaged by ID DPI” dataset, which contains all the 320 averaged observations. The averaged data were used to find the directions of maximal variance between individuals. A covariance matrix was generated and a PCA performed. PC1 accounted for 64.2 % of total variance and the first 4 PCs summed up to 87.4 % of it. PCs 4 and beyond accounted for less than 5 % of variation each and are therefore of little biological interest. Afterwards, PCA output was used for which is one of its main purposes in GM: visualization of shape change. Three types of graphs were obtained: PC shape changes (a diagram showing the shape changes associated with the PCs); Eigenvalues (a histogram showing the percentages of total variance for which the PCs account) and PC scores (a scatterplot of PC scores).

PC Scatterplots show the distribution of specimens along the axes of maximum variance (Figure 3A, B). To aid visualization, dots corresponding to early stages (3-6 DPI) were lightly colored and later ones (7-12 DPI) were darker colored. Results evidenced that PC1 is a development-related axis, because clearly separated early (mostly negative values) from late (positive values) stages of the experiment (Figure 3A). Moreover, at later stages ORMV-infected plants scored less positive in this axis, suggesting that infected plants retained a more juvenile (pedomorphic) shape. Positive extremes of PC2-4 are related to ORMV shapes.

**Figure 3.**
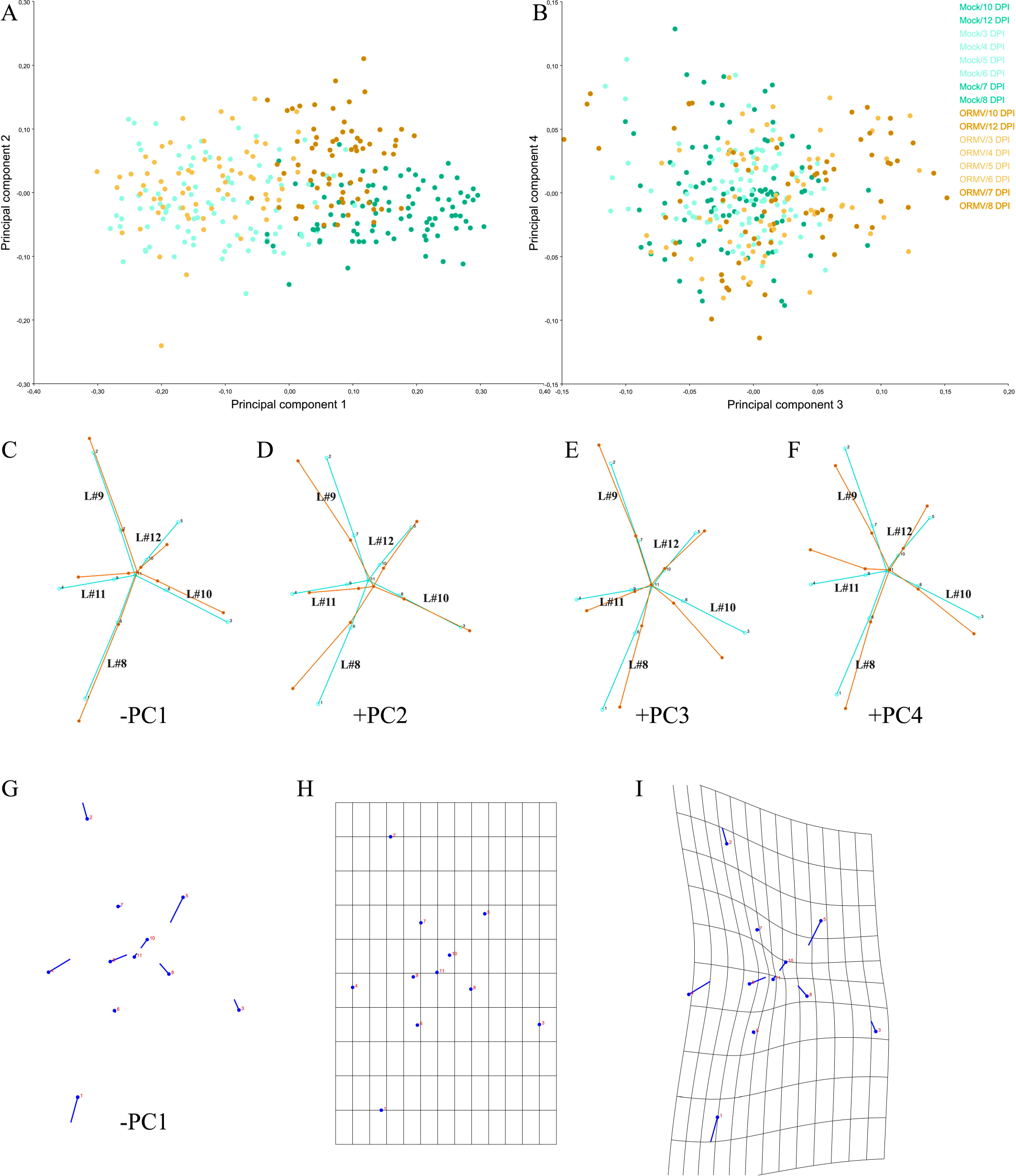
Shape variation between specimens (averaged by measurement replicates). PCA scatterplots of (A) PC1 vs. PC2 and (B) PC3 vs. PC4, which together explain 87.4 % of variance. Pale dots = juvenile (3-6 DPI) plants. Dark dots = mature (7-12 DPI) plants. (C-F) Wireframe graphs showing shape changes from the starting (average) shape (bluish green) to the target shape (orange) for the first four PCs. Negative (PC1) and positive (PCs2-4) components are shown, respectively. Here and throughout this work, leaf number is indicated in the wireframe in black. (G) Lollipop graph for the −PC1 component. Lollipops indicate starting position of landmarks with dots. (H-I) Transformation grids for (H) the starting shape and for (I) the target shape (-PC1). Shape changes (C-G and I) are magnified 2X for better visualization.

To this point, GM visualization tools are used to better understand what these relative positions on scatterplots mean respect to shape differences.

#### Visualization of shape changes

Prior to showing graphs from the “PC shape changes” tab, a brief description of common GM visualization tools is needed in order to accurately interpret the results. After the GPA, every configuration in the sample is optimally aligned to the average configuration and nearly optimally aligned to every other configuration in the sample. GPA removed differences attributable to size, position and orientation from configurations. All differences that remain are shape variation. Accordingly, shape differences are found using the relative displacements of the landmarks from one shape to another shape nearby in shape space (Klingenberg, 2013). By superimposing a target shape to a starting shape and looking for the relative displacement of homologous landmarks from one shape to another, insights on how variation between shapes occurs can be obtained and hypotheses about the underlying mechanisms, proposed.

A key concept to bear in mind is that it is fundamentally wrong to consider landmarks displacements in an isolated manner (Klingenberg, 2013; Zelditch et al., 2012). This is because all the landmarks included in the GPA jointly determine the alignment of each configuration in relation to the mean shape. Then, variation in the position of each landmark after superimposition is relative to the positions of all other landmarks. Though a shift is shown at every landmark, these shifts are relative to all other landmarks. Lollipop and wireframe graphs are based on these assumptions (see below).

Shape variation could be depicted by means of transformation grids. Transformation grids are mathematically constructed following the thin-plate spline technique, whose detailed explanation is far from the objective of this work and has been explained elsewhere (Klingenberg, 2013; Zelditch et al., 2012). Briefly, landmarks of a starting shape are placed on a grid of an imaginary infinitely thin metal plate. Landmarks of a target configuration are placed on another grid of equal characteristics, and both metal sheets are superimposed. Each landmark in the starting shape (e.g., mean shape) is linked to its homologous to reach the target configuration, and the deformation caused in the spline is calculated finding the smoothest interpolating function that estimates energy changes in the spline between landmarks. Importantly, differently from lollipop or wireframe graphs, transformation grids distribute the change in landmark positions to the space between landmarks, were no objective information is available. Then, whereas a powerful descriptive tool, transformation grids must be carefully interpreted, especially regarding regions of the object that do not have landmarks nearly positioned (Klingenberg, 2013; Zelditch et al., 2012,). More details and examples are given below.

Wireframe graphs (Figure 3C-F) connect the landmarks with straight lines for the starting and target shapes, thus showing the relative displacements of landmarks from a mean shape. Negative values of PC1 mostly correspond to juvenile (and infected) shapes; positive values of PC1 belong to healthy controls and adults. Hence, by depicting the −PC1 component, target shapes have negative values (Figure 3C). It can be seen that −PC1 explains the relative shortening of leaves #11 (landmarks 4 and 9) and #12 (landmarks 5 and 10). It makes sense, since younger plants have yet to develop these relatively new leaves. Petioles of leaves #10, #11 and #12 are particularly shortened. Relative to these shortenings, older leaves (#8 and #9) are relatively longer but, interestingly, only its laminae, since its petioles are not relatively elongated. Taken together, PC1 reveals that ORMV impaired the elongation of newer leaves to their normal extent. PC2 (Figure 3D) associates with relative radial displacements of leaves; tips of leaves #8 and #9 (landmarks 1 and 2) come close together, lowering the typical angle between successive leaves from near 137.5° to close to 90°. These relative displacements determine that leaves #9 and #10 form an exaggerated angle of near 180°. PC3 (Figure 3E) also mostly relates to radial changes in the infected rosette: leaf #10 is relatively displaced towards leaf #8 and the main effect is, again, the increase of the angle between leaves #9 and #10 to near 180°. PC4 (Figure 3F) explains less proportion of total variance (4.5%) and its effect is less clear; it is mostly related to the relative displacement of the lamina of leaf #11 towards leaf #9, almost without altering its petiole, which functions as a hinge. Leaves #9 and #10 are, as a combination of the effects depicted by PC2 and PC3, both relatively displaced towards leaf #8. Taken together, the wireframe visualization of the first four PCs (which account for more than 87% of total variance) show that ORMV induces the relative shortening of laminae and (especially) petioles of newest leaves, which relates to a pedomorphic shape, and provokes the relative displacement of leaves #9 and #10 towards leaf #8.

Displacement vectors (called “lollipop graphs” in MorphoJ) are arrows drawn between a landmark in a starting shape and the same landmark in a target shape. The dot in the lollipop represents the starting position and the vector is represented by a line departing from it (but in some software the inverse convention is followed, i. e., PAST). Though these visualization are being displaced in the GM literature in favor of more advanced tools (Klingenberg, 2013), here it is presented the case for −PC1, showing the relative displacements of landmarks (Figure 3G). It can be directly compared with Figure 3C.

Lastly, transformation grids are exemplified for −PC1 in Figure 3H-I. Figure 3H depicts the starting (mean) shape. Figure 3I show the transformed grid for −PC1. The compression of the grid in the central zone is the result of the relative displacement of landmarks 3, 8 (leaf #10), 4, 9 (leaf #11) and 5, 10 (leaf #12) towards the center of the rosette, whereas grid stretching is detected around landmarks 1 and 2 (revealing the relative expansion of laminae of leaves #8 and #9, since its petioles remain relatively immobile, landmarks 6 and 7). As stated above, visualization with these grids should be cautiously interpreted since the interpolation function deforms the grid between places where no landmarks are placed (and no information about even the existence of tissue is guaranteed). Therefore, only regions near landmarks should be discussed when viewing these graphs. To see these changes in more detail, PCA analyses were performed for each DPI. The “Combined dataset 3-12 DPI, averaged by ID DPI” was subdivided by DPI performing a common Procrustes fit, creating eight new datasets (DPIs) (raw data in Supplementary file ORMV.morphoJ). Covariance matrices were generated and a PCA performed for each DPI dataset. PC1 accounted for between 27-43 % of total variance and the first 4 PCs summed up from 78 to 84 % of it. PCs beyond PC4 accounted for 5 % or less of variation each. Shape change visualization showed that PC1 gradually separated specimens belonging to different treatments. Mock-inoculated plants were progressively more aligned with positive PC1 values. PC2 was more generally related to ORMV-infected plants in its positive values. Relative shortening of younger leaves and petioles, and relative displacement of leaves towards leaf #8 were progressively more accentuated (Supplementary Figure 2).

#### Discriminant Analysis

So far, differences between individuals were addressed with the aid of PCA. Afterwards a Discriminant Analysis (DA) was performed to test whether differences between treatments exist.

DA is a technique mathematically related to PCA. It finds the axes that optimize between-group differences relative to within group variation. It can be used as a diagnostic tool (Zelditch et al., 2012). It is here used for testing treatments using a series of tests for sample mean differences including an estimate of the accuracy of shape in predicting groups. The capability of DA to correctly assign specimens to treatments was assessed along the experiment using the averaged datasets for each DPI. In MorphoJ, Discriminant Function Analysis was requested selecting “Treatment” as classification criterion. By default, DA in MorphoJ performs a parametric Hotelling’s T-square test, and here there were also requested permutations tests for the Procrustes and Mahalanobis distances with 1000 random runs. Hotelling’s test is the multivariate equivalent of the common Student´s t-test. Procrustes and Mahalanobis distances show how far shapes from one group are from the mean of the other group. Results of the tests are shown in Table 1. At 5 DPI the three tests found shape differences between treatments (0.001<p<0.005). From 6 DPI and beyond, p-values were extremely significant (p<0.0001). These results coincide with those obtained by Procrustes ANOVA of shape (Supplementary Table 2). DA maximizes group separation for plotting their differences and predicting group affiliation (classification). The classification of a given specimen (through the discriminant axis) is done using functions that were calculated on samples that included that same specimen (resubstituting rate of assignment). Then, a degree of over-fitting is unavoidable and leads to an overestimate of the effectiveness of the DA. To overcome this problem, one solution is to use a cross-validation or jackknife procedure (Viscosi and Cardini, 2011; Zelditch et al., 2012). Jackknife procedure leaves one specimen at a time not used for constructing the Discriminant function and then tests the rate of correct specimen assignment. Only jackknife cross-validated classification tables provide reliable information on groups. Results of DA in group assignment are shown in Figure 4 for 3, 7 and 12 DPI and detailed for all DPIs in Supplementary Table 3. As expected, resubstitution rates of assignment (Figure 4A, D, G) were higher than jackknifed counterparts (Figure 4B, E, H), but the latter reached high levels of accuracy (≥90 %) from 6 DPI and beyond (Supplementary Table 3). To test whether this level of accuracy was indeed good, these results were compared with classification/misclassification tables completed by human observers. The entire image dataset of 7 DPI was given to three expert researchers working with Arabidopsis (one of the authors (S. Asurmendi) and two other researchers from another Institution). They were all blind to the assignment of treatments to each plant, except for one Mock-inoculated and one ORMV-infected plant that were given as references. They classified the 38 remnant plants and results are shown in Supplementary Table 3. Human accuracy ranged from 55 to 72.5 %, with an average of 64.2 %. Therefore, DA outperformed expert human eye by 30 % at 7 DPI and yielded higher classification rates from 5 DPI.

**Figure 4.**
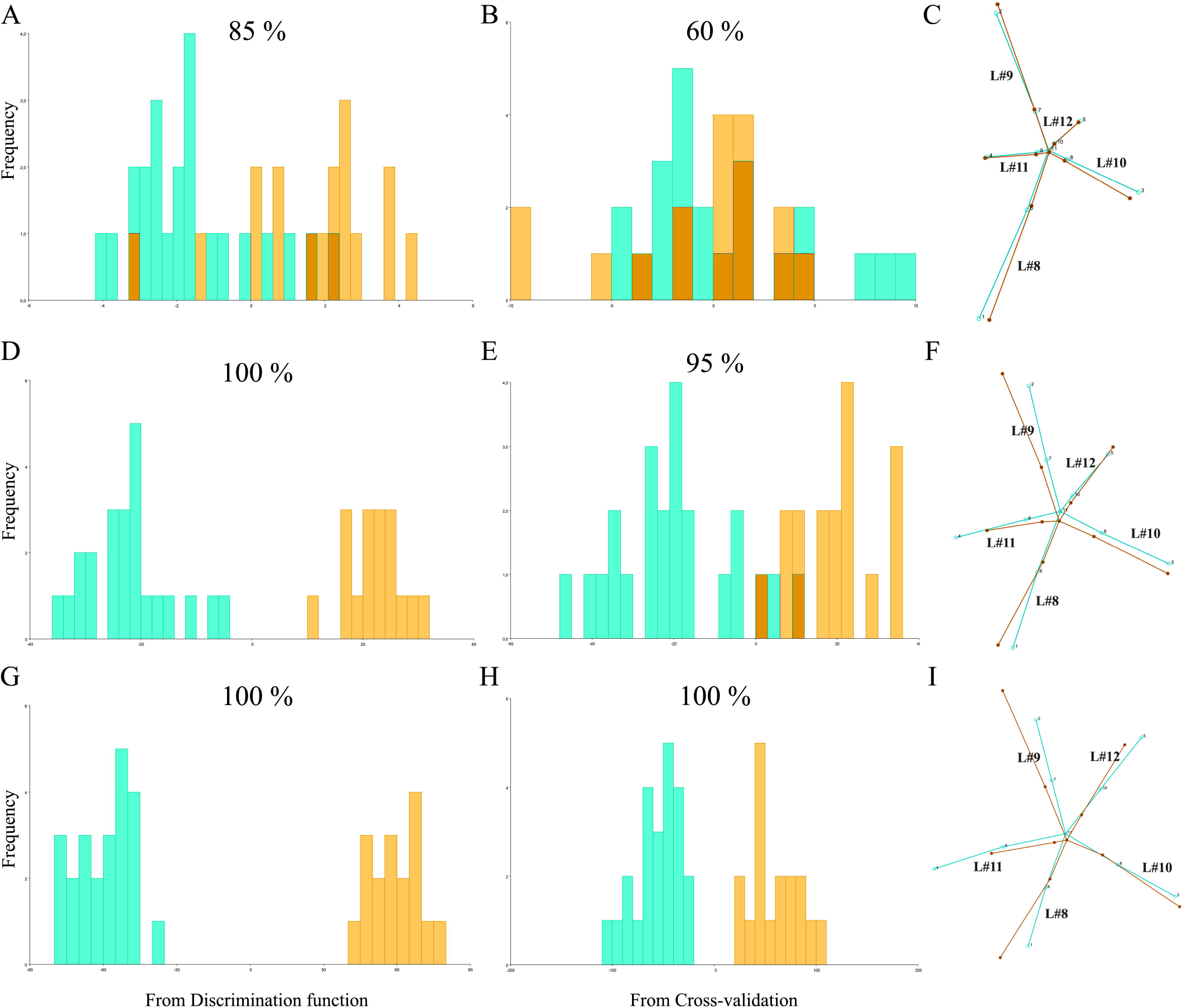
Discriminant analyses of shape variation between treatments at 3 (A-C), 7 (D-F) and 12 (G-I) DPI. Frequencies of discriminant scores obtained by resubstitution rates of assignments (A, D, G) and a jackknife cross-validation (B, E, H) are shown using histogram bars with percentages of correct assignments above each graph. Wireframes comparing mean shapes (C, F, I) are shown magnified 2 times. Mock = bluish green; ORMV = orange.

**Table 1.**
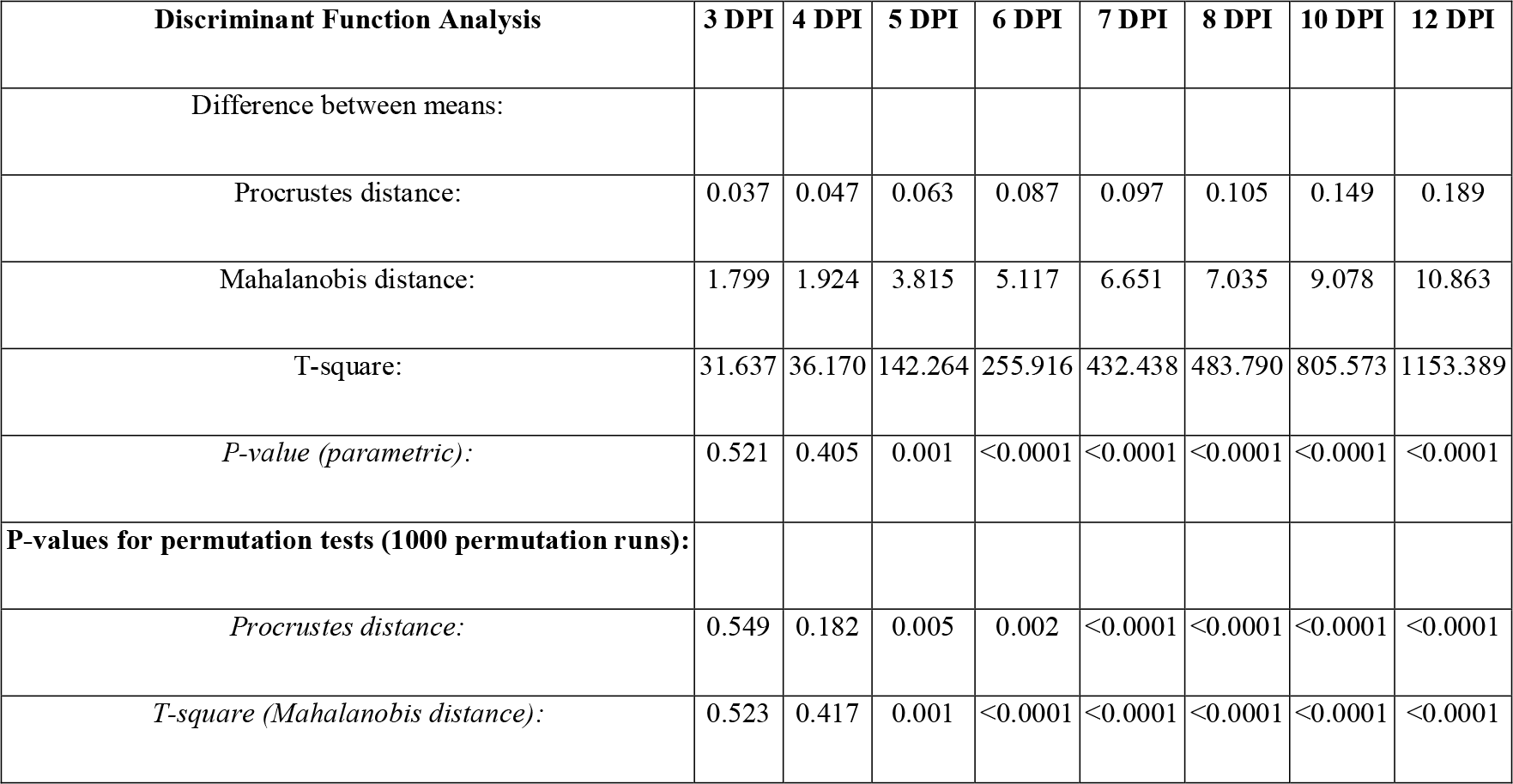
Statistical tests for differences between means of treatments at each DPI from DA. Permutation tests with 1000 random runs.

| **Discriminant Function Analysis** | **3 DPI** | **4 DPI** | **5 DPI** | **6 DPI** | **7 DPI** | **8 DPI** | **10 DPI** | **12 DPI** |
| --- | --- | --- | --- | --- | --- | --- | --- | --- |
| Difference between means: |  |  |  |  |  |  |  |  |
| Procrustes distance: | 0.037 | 0.047 | 0.063 | 0.087 | 0.097 | 0.105 | 0.149 | 0.189 |
| Mahalanobis distance: | 1.799 | 1.924 | 3.815 | 5.117 | 6.651 | 7.035 | 9.078 | 10.863 |
| T-square: | 31.637 | 36.170 | 142.264 | 255.916 | 432.438 | 483.790 | 805.573 | 1153.389 |
| *P-value (parametric):* | 0.521 | 0.405 | 0.001 | <0.0001 | <0.0001 | <0.0001 | <0.0001 | <0.0001 |
| **P-values for permutation tests (1000 permutation runs):** |  |  |  |  |  |  |  |  |
| *Procrustes distance:* | 0.549 | 0.182 | 0.005 | 0.002 | <0.0001 | <0.0001 | <0.0001 | <0.0001 |
| *T-square (Mahalanobis distance):* | 0.523 | 0.417 | 0.001 | <0.0001 | <0.0001 | <0.0001 | <0.0001 | <0.0001 |

Wireframe graphs for 3, 7 and 12 DPI (Figure 4C, F, I) show the difference from Mock group to ORMV group. There is little difference at 3 DPI, if any (Figure 4C), consistently with nonsignificant differences found by DA at this stage. At 7 DPI (Figure 4F), the relative shortening of leaf #11 (landmarks 4 and 9) is evident, as is the relative increase in the angle between leaves #9 and #10. These tendencies persisted at 12 DPI (Figure 4I). At this stage, petioles of leaves #11 and #12 are strongly relatively shortened. These results resemble those obtained in Figure 3C-F and approximately summarize shape changes explained by the first 4 PCs, indicating that these shape differences not only separated juveniles from adults but are also hallmarks of shape change induced by ORMV. These results are interesting because discriminant axes not necessarily resemble PCA axes (Zelditch et al., 2012).

### Allometric patterns and size correction

As ORMV induced not only changes in shape, but also in size (Supplementary Table 2) it is worth investigating whether shape changes are associated to size differences. In principle, group differences could arise if individuals of one group are different in shape because they grew faster than the other group´s individuals and reached earlier a more advanced developmental stage. The association between a size variable and the corresponding shape variables is called allometry. Isometry, by contrast, is the condition where size and shape are independent of each other and usually serves as the hypothesis null. These concepts are rooted in the Gould-Mosimann school of allometry that conceptually separates size and shape (Klingenberg, 2016). Though size had been removed from forms after GPA, leaving shape differences free of it, there could be a linear relationship between them. Allometry can be statistically tested for by tests of multiple correlation.

When groups are present, a single regression line through all groups cannot be fit to test allometry because lines could have group-specific slopes or intercepts (Viscosi and Cardini, 2011). As TPSRegr (see below) uses raw data coordinates and averaged by ID DPI datasets in MorpohoJ do not have them, these analyses were carried on with individual datasets from only the 1^st^ Digitization. As proven earlier, differences between digitizations were negligible (Figure 2 and Supplementary Table 2).

To test whether an allometric component is present in each group, separate regressions were performed for each treatment and DPI with Procrustes Coordinates as dependent variables and ln(CS) as the independent variable. Permutation tests were requested with 10.000 runs. Respective p-values and predicted SS from regressions (which correspond to allometric variation of shape) are shown in Figure 5A. Allometry accounted for moderate to high proportions of the total shape variation since SS reached values of 36% at 6 DPI (Mock). ORMV induced a reduction in the allometric component of shape variation as evidenced by lower predicted SSs along the experiment and non-significant values of allometry for all except 4 and 5 DPIs. For both treatments and particularly for healthy controls, a bell-shaped curve is detected and a maximum of allometry is seen at 6 DPI for Mock plants but a day before for ORMV. Differences between treatments start sharply at 5 DPI, when allometry accounts for 32% of predicted SS for Mock but only 20% for ORMV. This analysis shows that for ORMV, shape variation is much less driven by size heterogeneity (at a given DPI) and that for Mock plants this situation (isometry) occurs at later stages of development (10-12 DPI).

**Figure 5.**
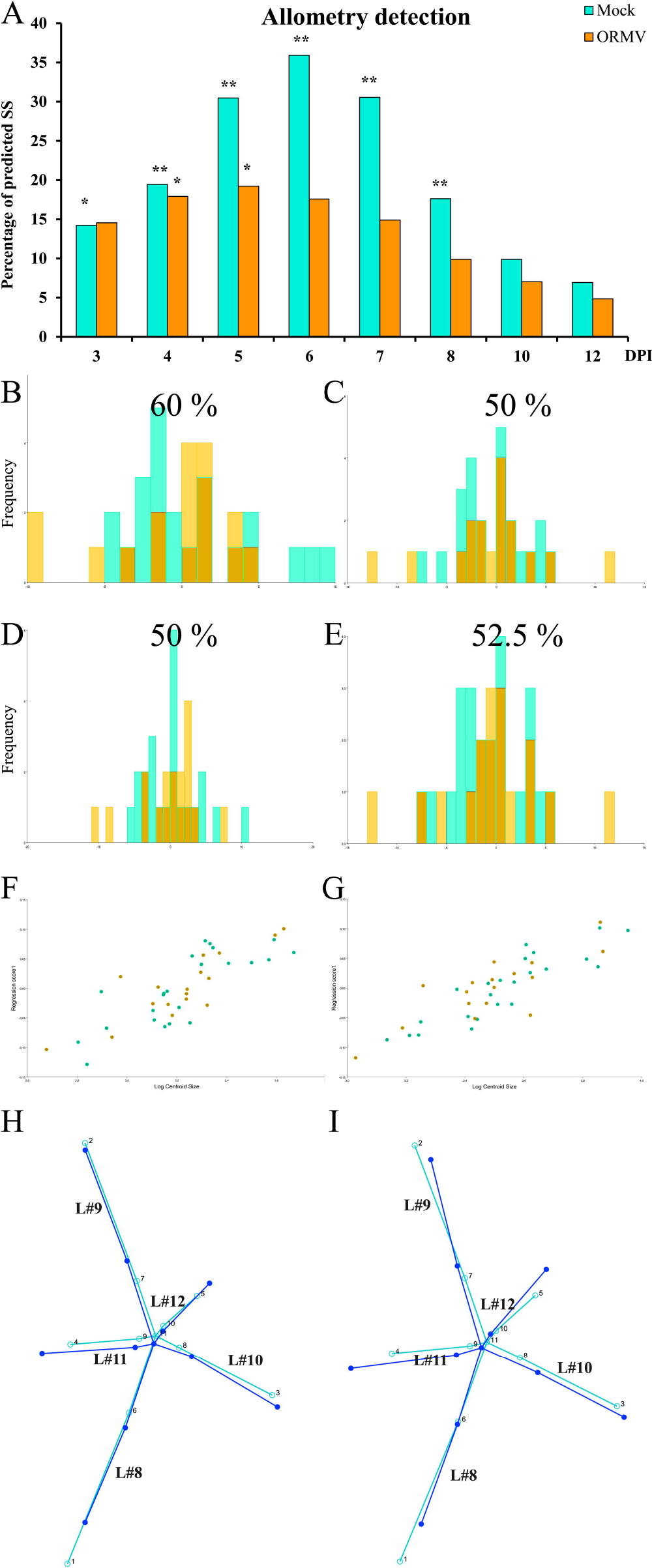
Allometric analyses. (A) Predicted sum of squares from regressions of shape onto ln(CS) for each treatment and DPI. P-values were corrected using Holm´s sequential test (*α*=0.05). * = p < 0.05; ** = p < 0.01. Allometric analyses for (B, D, F, H) 3 DPI and (C, E, G, I) 4 DPI (Mock = bluish green; ORMV = orange). Cross-validated DAs before (B-C) and after (D-E) size correction with percentages of correct assignments above each graph. (F-G) Scatterplot of regression scores vs. ln (CS). (H-I) Wireframes showing starting mean shape (turquoise) and target shape depicting an increase in one unit of ln(CS) (blue), without magnification.

When at least one group has regression slopes different from zero a series of tests could be done in order to control for size and repeat analyses to assess whether differences in shape are actually the result of size variation only (Klingenberg, 2016; Viscosi, 2015; Viscosi and Cardini, 2011; Zelditch et al., 2012). TPSRegr (v. 1.41) was used firstly to determine whether group-specific slopes were parallel at each DPI (3-8). Only at 3 and 4 DPI this occurred (p > 0.05, slopes not statistically significant). As slopes were found to be parallel, it is possible to test whether they are separate parallel slopes or coincident (same Y-intercept). TPSRegr tests demonstrated that slopes are coincident (p > 0.05). Then, size-corrections could only be done for 3 and 4 DPI, since from 5 to 8 DPI slopes were different (p < 0.05) and groups follow its own allometric pattern and for 10 and 12 DPI there are isometry and size do not correlate with shape variation. Size-correction was done for 3 and 4 DPI separately in MorphoJ, using all 40 plants. Shape variables were regressed onto ln(CS) for each dataset, pooling regressions within subgroups (treatments) and permutation tests with 10.000 runs were requested. Residuals from the analyses contain the size-free information about shape only and can be used to repeat DAs to test for improved accuracy of discrimination (Klingenberg, 2016). Results (Figure 5B-E) showed that group separation was not improved. This is somewhat expected since at this stage of viral infection there are no detectable differences in size nor shape yet (Figure 4 and Tables 1-3). This test and the large overlap between populations in the scatterplot of regression scores onto size (Figure 5F, G) suggest that the effect of size on shape is very similar for both treatments and DPIs: bigger rosettes have further distal displacements of leaves #10, 11 and 12 relative to older leaves (#8 and #9) and elongated petioles (Figure 5H, I) thus reflecting the differential internal growth of the rosette. Bigger, more mature rosettes have more developed newest leaves.

### Phenotypic Trajectory Analyses (PTA)

Whilst the comparison of allometric vectors indicated that shape change is altered at definite DPIs during ORMV infection, a holistic view of ontogenetic alterations needs to measure phenotypic evolution across multiple levels. It allows ontogenetic patterns to be characterized as phenotypic trajectories through the morphospace, rather than phenotypic vectors. The method proposed by (Adams and Collyer, 2009) “*(…) may also be used for determining how allometric or ontogenetic growth trajectories differ, or for quantifying patterns in other data that form a time-sequence*” (Adams and Collyer, 2009). Briefly, phenotypic trajectories have three attributes: size, direction and shape.

Trajectory size (*MD*) quantifies the path length of the phenotypic trajectory expressed by a particular group across levels. This represents the magnitude of phenotypic change displayed by that group. If trajectories of two or more groups compared over comparable time periods differ in trajectory size then it indicates differences in rates of morphological change.

Trajectory direction (*θ*) is a multivariate angle that describes the general orientation of phenotypic evolution in the multivariate trait space. Statistical comparisons of trajectory direction can be used to provide an assessment of patterns of convergence, divergence, and parallelism.

Trajectory shape (*D*_*Shape*_) describes the shape of the path of phenotypic evolution through the multivariate trait space. This information is useful because it indicates whether there are differences in how each group occupies the morphospace through the time period.

PTA analysis proceeds by starting from the PCs for all specimens at all DPIs. They were obtained from the “Combined dataset 3-12 DPI, averaged by ID DPI” of the Supplementary file ORMV.morphoj. The R script developed by (Adams and Collyer, 2009) was run in RStudio (RStudio Team, 2016).

PTA approach (with 1,000 residual randomization permutations) revealed significant differences in the magnitude of phenotypic evolution between the two treatments (*MD*_Mock,ORMV_ = 0.100, P_*size*_ = 0.003), implying that ORMV-infected plants experienced a lower rate of ontogenetic phenotypic evolution relative to controls. Overall direction of ontogenetic changes were also statistically significantly different (*θ*_Mock,ORMV_ = 18.34°, P_*θ*_ = 0.001). Finally, shape assessment analysis showed differences between treatments regarding trajectories over time (*D*_*Shape*Mock, ORMV_ = 0.367, P_Shape_ = 0.001) (Table 2). When phenotypic trajectories are plotted through time for the first two principal components (Figure 6) these statistical conclusions are graphically confirmed. Group trajectories diverge from 3 DPI and trajectory lengths are evidently different, specifically regarding the relative stasis of the ORMV-infected group beyond 6 DPI. Both factors contribute to the overall difference found in trajectory shape.

**Figure 6.**
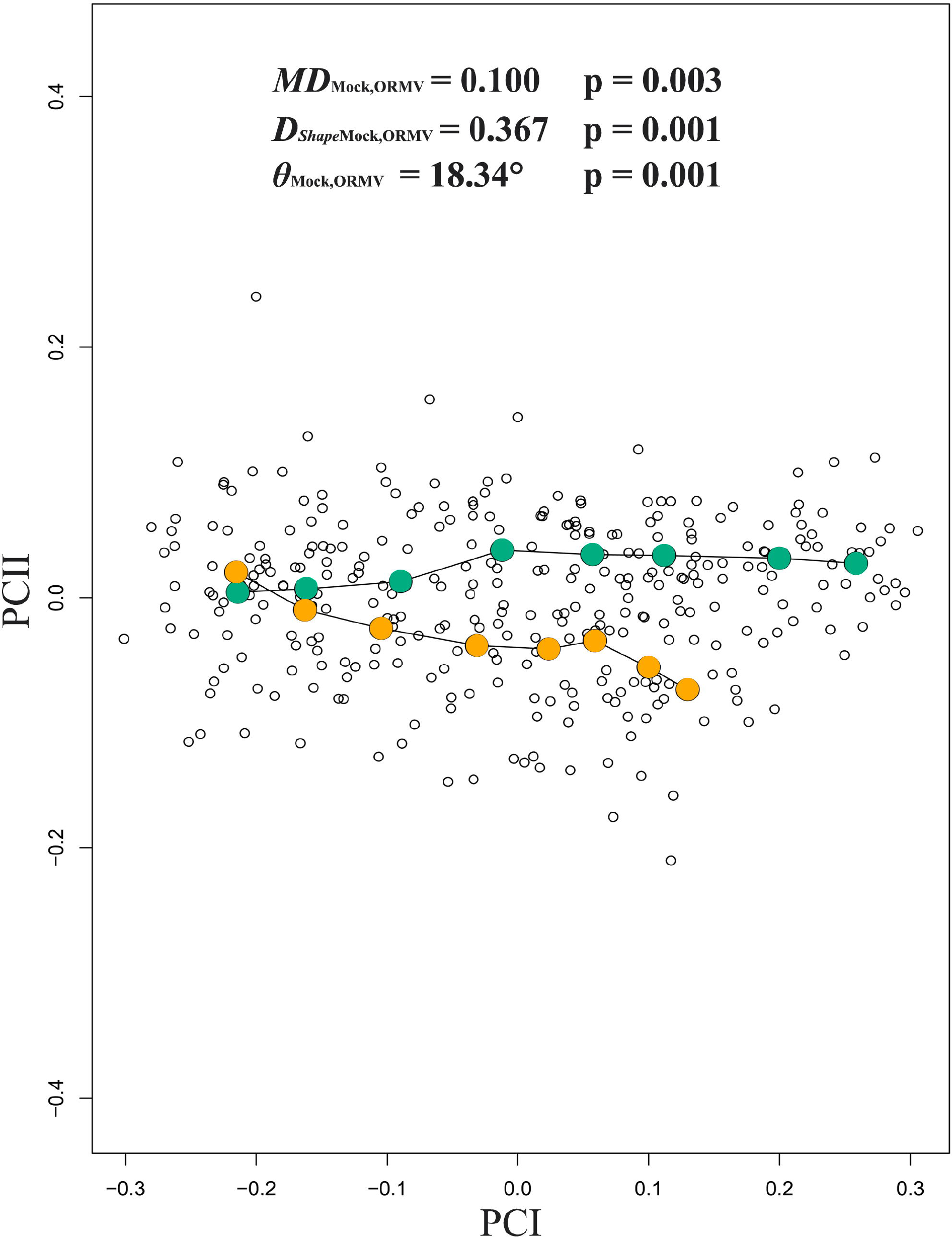
Phenotypic trajectories for Mock and ORMV (3-12 DPI). Scatterplot shows the first two PCs of shape variation across the experiment. Mean values for each DPI are colored and connected with lines. PTA parameters are given (see main text). Mock = bluish green; ORMV = orange.

**Table 2.** Comparative trajectory analyses for the full dataset of the ORMV experiment (3-12 DPI), the reduced dataset (4-10 DPI) and the comparisons with TuMV experiments (4-10 DPI).

| ORMV3-12 DPI |  |  |
| --- | --- | --- |
|  |  | *p-value* |
| $MD_{Mock,ORMV}$ | 0.100 | **0.003** |
| $\theta_{Mock,ORMV}$ | 18,34° | **0.001** |
| $D_{ShapeMock, ORMV}$ | 0.367 | **0.001** |
| $MPL_{Mock}$ | 0.696 | **2.50E-04** |
| $MPL_{ORMV}$ | 0.596 | |
| $\sum Var_{Mock}$ | 0.035 | **2.52E-06** |
| $\sum Var_{ORMV}$ | 0.023 | |
| $D_{1(Mock)}$ | 20.21 | **2.89E-06** |
| $D_{1(ORMV)}$ | 26.93 | |
| Hyperellipse<sub>(IC95%)Mock</sub> | 0.022 | **0,005 \*** |
| Hyperellipse<sub>(IC95%)ORMV</sub> | 0.014 |  |
| $D_{2(Mock)}$ | 32.97 | **0,040 \*** |
| $D_{2(ORMV)}$ | 41.87 | |
| ORMV4-10 DPI |  |  |
| $MD_{Mock,ORMV}$ | 0.085 | **0.005** |
| $\theta_{Mock,ORMV}$ | 16,46° | **0.001** |
| $D_{ShapeMock, ORMV}$ | 0.343 | **0.037** |
| $MPL_{Mock}$ | 0.472 | **8,03E-04 \*** |
| $MPL_{ORMV}$ | 0.401 | |
| $\sum Var_{Mock}$ | 0.022 | **2,21E-05 \*** |
| $\sum Var_{ORMV}$ | 0.015 | |
| $D_{1(Mock)}$ | 21.77 | **3,61E-05 \*** |
| $D_{1(ORMV)}$ | 28.37 | |
| Hyperellipse<sub>(IC95%)Mock</sub> | 0.012 | 0,075 * |
| Hyperellipse<sub>(IC95%)ORMV</sub> | 0.009 |  |
| $D_{2(Mock)}$ | 48.94 | 0,203 * |
| $D_{2(ORMV)}$ | 56.41 | |
| TuMV4-10 DPI 1st |  |  |
| $MD_{Mock,TuMV}$ | 0.093 | **0.015** |
| $\theta_{Mock,TuMV}$ | 34,41° | **0.001** |
| $D_{ShapeMock, TuMV}$ | 0.613 | **0.001** |
| $MPL_{Mock}$ | 0.504 | **0,049 \*** |
| $MPL_{TuMV}$ | 0.461 | |
| $\sum Var_{Mock}$ | 0.030 | **0,007 \*** |
| $\sum Var_{TuMV}$ | 0.023 | |
| $D_{1(Mock)}$ | 16.94 | **0,017 \*** |
| $D_{1(TuMV)}$ | 21.83 | |
| Hyperellipse<sub>(IC95%)Mock</sub> | 0.019 | 0,156 * |
| Hyperellipse<sub>(IC95%)TuMV</sub> | 0.017 |  |
| $D_{2(Mock)}$ | 32.51 | 0,277 * |
| $D_{2(TuMV)}$ | 46.05 | |
| TuMV4-10 DPI 2nd |  |  |
| $MD_{Mock,TuMV}$ | 0.082 | 0.202 |
| $\theta_{Mock,TuMV}$ | 35,05° | **0.001** |
| $D_{ShapeMock, TuMV}$ | 0.642 | **0.002** |

\*= First 3 PCs considered (>95% total variance)
Units: MD = DShape = MPL = D1 = D2 = Euclidean distance.
$\theta$ = degrees. $\sum Var$ = Hyperellipse(CI=95%) = dimensionless.
Statistically significant results in bold

However, as pointed out by (Ciampaglio et al., 2001), no one method of disparity measurement is sufficient for all purposes. Using a combination of techniques should allow a clearer picture of disparity to emerge. With this aim, another available approach to compare shape trajectories through multivariate morphospace was used. Originally developed to study unequal morphological diversification in a clade of South American fishes (Sidlauskas, 2008), this approach is useful because allows investigating whether a group “explores” different amount of morphospace than others, additionally to possible differences in magnitude of phenotypic evolution. Moreover, density parameters (D) could be calculated to determine whether the amount of morphological change is more or less constrained in the morphospace.

The method was adapted to the present case study: as there is not a phylomorphospace and both treatments lack a “common ancestor” but each plant follow its own independent ontogenetic path, nodes and branches do not exist. Rather, each plant possesses its own trajectory without points in common. Taken these considerations into account, morphological trajectories were calculated for all plants. To do so, the “Combined dataset 3-12 DPI, averaged by ID DPI” of the Supplementary file ORMV.morphoj was subdivided by ID. Forty new datasets (Mock- and ORMV-inoculated plants from the same previously performed Procrustes fit) were obtained and Procrustes Coordinates and eigenvalues from the 7 PCs obtained were exported to an Excel spreadsheet.

The morphometric change experienced by a plant throughout ontogeny equals the Euclidean distance (D) between successive points in morphospace that represent its shape at each DPI. As PCs from a PCA carry all the morphological information extracted from the Procrustes Coordinates, distances are simultaneously calculated over all the PCs using the Pythagorean Theorem. These distances are designated as morphometric path lengths (∑D = MPL) (sensu (Sidlauskas, 2008)). Mock-inoculated plants traveled on average more distance through morphospace than infected ones (MPL_Mock_ = 0.6956 vs. MPL_ORMV_ = 0.5963, p = 0.00025, Mann-Whitney test). Other measures are traditionally used to detect changes in morphospace occupation patterns and the amount of difference between character states among specimens in morphospace (Ciampaglio et al., 2001), e.g. sum of variances (∑Var). Control plants had higher ∑Var values than infected plants (∑Var _Mock_ = 0.0350 vs. ∑Var_ORMV_ = 0.0230, p = 2.52 x 10^−6^, Mann-Whitney test) a result that pointed to a higher increase in shape change in controls (Ciampaglio et al., 2001). Morphospace density occupation measures could be obtained taking into accounts not only MPLs but also variances of the PCs across the experiment. If a group folded an equivalent amount of morphometric change into a much smaller region of morphospace than the other, thus will have a higher density (Sidlauskas, 2008). Morphometric path density (D) could be calculated as D_1_ = MPL/∑Var. ORMV-infected plants are more densely restricted in morphospace (D_1(Mock)_ = 20.21 vs. D_1(ORMV)_ = 26.93, p = 2.89 × 10^−6^, Mann-Whitney test) (Table 2).

An alternative measure of density (D_2_ = MPL/V) considers the volume (V) that the group occupies in morphospace. A variety of volumetric measures are possible (Ciampaglio et al., 2001). This study considered the volume of a 95% confidence hyperellipse. It was obtained by calculating the square root of the product of the eigenvalues of the PCs and comparing them with expected values for a X^2^ distribution at α = 0.05 (see Materials and Methods). Mock-inoculated plants have hyperellipses of higher volume on average than infected plants (Hyperellipse_(IC95%)Mock_ = 0.0129 vs. Hyperellipse_(IC95%)ORMV_ = 0.0073, but differences were not statistically significant (p = 0.11888, Mann-Whitney test). Similarly, density measures based on hyperellipses calculations were not statistically significantly different (D_2(Mock)_ = 111.47 vs. D_2(ORMV)_ = 146.34, p = 0.25051, Mann-Whitney test), although ORMV-infected plants had a higher average density. These differences in density measures could arise from the fact that hypervolume calculations can produce values that are extremely small and variable. Since the hypervolume is calculated by taking the product of univariate variances, any axis or axes with negligible variance will produce a value of hypervolume close to zero. Moreover, all multiplied variances are given the same weight and consequently, PC axes that represent a minimal percentage of the total variance could distort conclusions obtained with more informative axes. Thus, hypervolume can be very sensitive to variation in a single character. To avoid this issue, only the axes with significant variances are chosen to represent the disparity among points in morphospace (Ciampaglio et al., 2001). Therefore, the analysis was repeated including only the first three PCs, which accounted for more than 95% of variance. Results were similar to previously obtained for all parameters but hyperellipse´s volumes were found to be statistically significantly different (Hyperellipse _(IC95%)Mock_ = 0.022 vs. Hyperellipse_(IC95%)ORMV_ = 0.014, p = 0.0052597, Mann-Whitney test) as the D_2_ parameter (D_2(Mock)_ = 32.97 vs. D_2(ORMV)_ = 41.87, p = 0.040172, Mann-Whitney test) (Table 2).

Taken together, PTA showed that Mock-inoculated and ORMV-infected plants follow separate paths through morphospace. They differ in length, direction and shape (Figure 6), but also explore distinct regions of morphospace in a disparate quantity. Control plants experience more diversification of shape, as evidenced by the comparative length of trajectories (*MD* and MPL), have a higher amount of difference between shape states through the experiment in morphospace (∑Var) and explore more ample regions of morphospace (D_1_, D_2_) (Table 2). On the whole, ORMV infection not only alters the direction of ontogenetic shape development but also diminishes shape change.

### Growth and Development modeling

Even after finding that Mock- and ORMV-infected plants follow different ontogenetic trajectories of shape it is possible to compare their rates and timings of growth and development. When groups have different ontogenetic trajectories of shape, it is necessary to use a formalism that can be used when treatments follow group-specific ontogenetic trajectories (Zelditch et al., 2012). One such possibility is to compare the rates and timings at which groups depart from their own juvenile forms (Gould, 1977), an approach that can be applied to compare growth (Hingst-Zaher et al., 2000) and development (Zelditch et al., 2003) rates between groups with different ontogenetic trajectories. To linearize the relationship between size and age, ln(CS) was regressed on ln(DPI) and growth rates were compared. Results showed higher growth rate for Mock (1.06, CI_95%_=1.00-1.12) than for ORMV (0.72 CI_95%_=0.64-0.79) (p_(same slope)_= 3,0309E-12) (Figure 7A). Lack-of –fit was assessed for both regressions and rejected (p= 0.9975 and p= 0.3144 respectively), thus indicating the goodness of fit for both linear regressions. To compare developmental rates, it was measured the rate at which shape progressively differentiates away from that of the youngest age class (3 DPI) from 4 to 12 DPI. The degree of differentiation is measured by the morphometric distance between each individual and the average of the youngest age class (Zelditch et al., 2003), using Euclidean distances as approximations of Procrustes distances (Supplementary Figure 1 and Supplementary Table 1). Linear regressions with Euclidean distances (D) as a dependent variable and ln(DPI) as a regressor indicated a higher developmental rate for Mock- (0.34, CI_95%_=0.32-0.36) relative to ORMV-infected plants (0.24, CI_95%_=0.22-0.26) (p_(same slope)_ = 5,4657E-13) (Figure 7B). Lack-of –fit was rejected (p= 0.1626 and p= 0.3278 respectively). Healthy controls depart more from its own juvenile shape from 8 DPI and beyond (Figure 7C), indicating that developmental change was relatively impaired by ORMV. Together, these results indicated that ORMV reduced both growth and morphological change.

**Figure 7.**
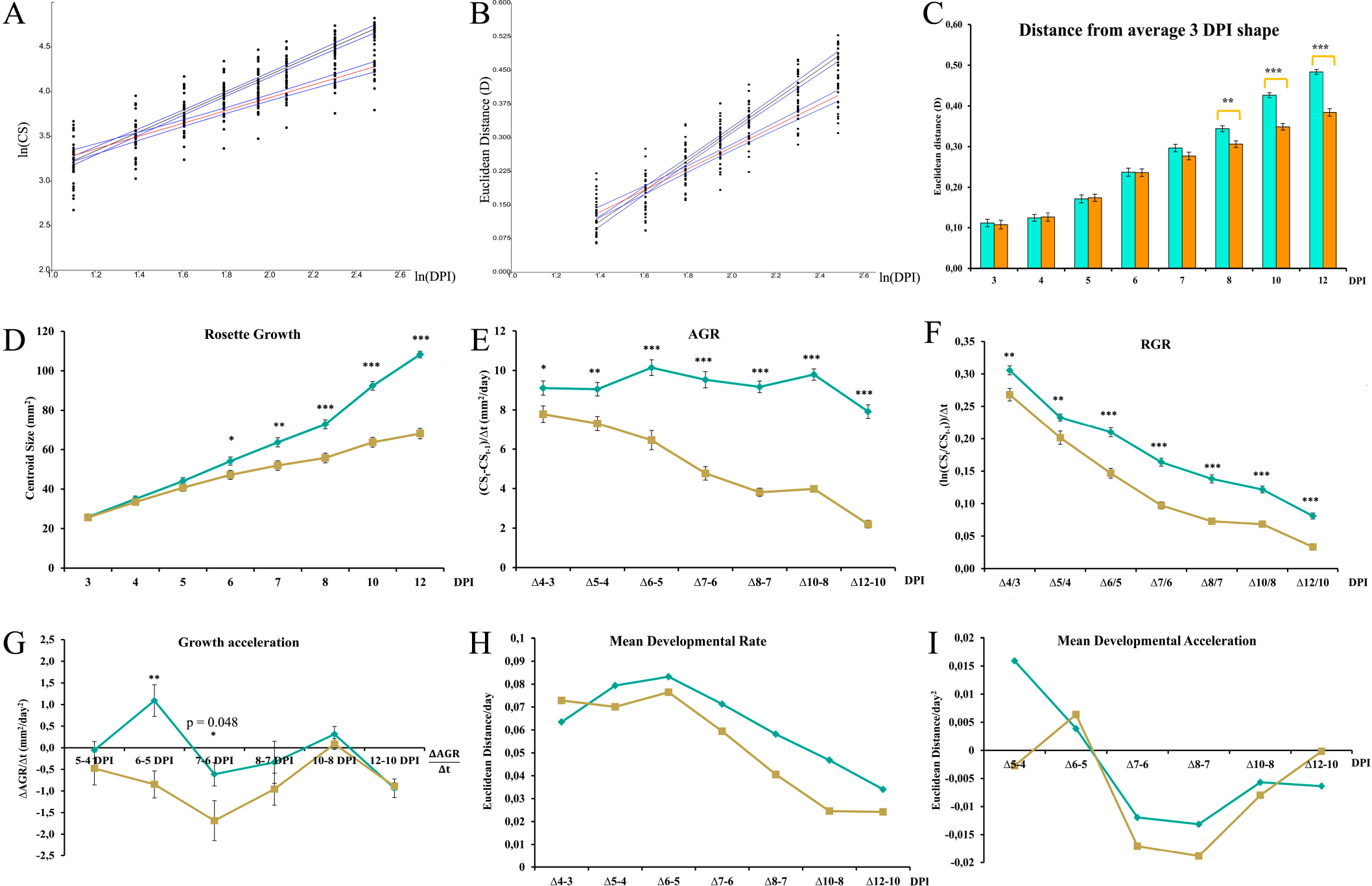
Growth and Development modeling. Comparisons of (A) growth and (B) developmental rates. Linear regressions for Mock (black lines) and ORMV (orange) with CI95% bands (blue). (C) Euclidean distances from own average juvenile shapes for mock and ORMV plants. Student´s t tests were performed separately for each DPI, contrasting mock vs. ORMV mean distances from its own average shapes at 3 DPI. Bars indicate mean average shape distances from average juvenile shape +/− SE. ** = p < 0.01; *** = p < 0.0001. (D-G) Rosette growth parameters. Measures of (D) size, (E-F) growth rate and (G) growth acceleration. Error bars indicate +/− SE. * = p < 0.05; ** = p < 0.01; *** = p < 0.0001, Mann-Whitney tests. (H-I) Rosette developmental parameters. (H) rate and (I) acceleration. (C-I) Mock = bluish green; ORMV = orange.

Alternatively, nonlinear models have been widely applied to studies of growth in several biological species (reviewed in (Zelditch et al., 2012)), including Arabidopsis (Tessmer et al., 2013). The latter decided to apply a logistic model regarding Arabidopsis growth from seedling stage on the basis of the prevalence of that model in plant growth studies. However for the present study it was decided to follow a less aprioristic approach, more in line with that of (Zelditch et al., 2003) who tested several nonlinear models and compared their relative performance regarding absence of residuals autocorrelation, percentage of variance explained and minimal parameterization. Here, five nonlinear models were chosen to compare: Logistic, Gompertz, Exponential, Monomolecular and Richards (Zelditch et al., 2003). For development analysis (Euclidean distances respect to 3 DPI mean shape) Logistic model fitted the best, with minimal Mean Square Error and low correlation between parameters, very close to the Gompertz model. However, all tested models showed a strong autocorrelation of residuals as evidenced for the Durbin-Watson panel test (Appendix S1). When the residuals are autocorrelated, it means that the current value is dependent of the previous (historic) values and that there is a definite unexplained pattern in the Y variable (Euclidean distance in this case study) that shows up in the disturbances. As a basic assumption of these analyses is the independence of residuals and particularly, their absence of autocorrelation, neither analyzed model fitted the development accurately. The same problem was found when the five growth models were applied to study growth, regardless choosing CS or ln (CS) as the dependent variable and DPI or ln(DPI) as the regressor (data not shown). An explanation for the autocorrelations of residuals is that data from successive DPIs (intra-group analyses) are not independent: every specimen is recorded at each DPI. This leads to a multivariate longitudinal data analysis situation, a branch of statistical analysis that has been recently addressed following different approaches (Verbeke et al., 2014), and whose level of complexity is beyond the scope of this work.

However, intra-treatment paired comparisons of shape are possible using a paired Hotelling´s test (a multivariate analog of the paired t-test). It was found a strong effect of time on shape and differences are extremely statistically significant for Mock plants (Supplementary Table 4).

Basic rosette growth and development parameters were studied (Figure 7). Rosette Growth, Absolute Growth Rate (AGR) and Relative Growth Rate (RGR) had been proposed as measures of rosette expansion and its velocity and rate, respectively (De Vylder et al., 2012; Vanhaeren et al., 2015). Mock-inoculated plants were statistically significantly bigger from 6 DPI and beyond (Figure 7D). The graphic lacks the typical sigmoidal shape of growth curves, probably because early stages of development (when landmarks used in this work were not present yet) were not included in the analysis and plant growth had not reached its plateau phase at 12 DPI yet. AGR and RGR analyses revealed an early change in growth tendencies between treatments. As early as between 3 and 4 DPI, (two days before the detection of significant differences in rosette area) ORMV started to slow rosette growth relative to healthy controls (Figure 7E-F). AGR graphic (Figure 7E) shows that, in contrast with control plants, ORMV-infected plants grew less rapidly from one day to the following throughout the experiment with the exception of the period between 8 and 10 DPI. This indicated different growth acceleration for each treatment. Growth acceleration (Figure 7G) peaked between 5 and 6 DPI in control plants and remained near zero until 12 DPI, suggesting a stage of linear rate of expansion. Later on, negative acceleration could indicate an entering in plateau phase reached by the region of the rosettes under study. In infected plants, however, acceleration between 5 and 6 DPI was negative and differed strongly from controls (difference = 1.94 mm^2^/day^2^, p= 0.0009, Mann-Whitney test), indicating that ORMV early slowed down the velocity of plant growth in a drastic manner. A trend towards more negative values of growth acceleration relative to controls was maintained in ORMV-infected plants until 10 DPI, though only marginally statistically significant. At 10-12 DPI both groups decelerate their growth, possibly indicating an entering in a plateau phase of growth. These results indicate that ORMV induces measurable changes in growth rates before the mean CS is found to be statistically significantly different from healthy controls, and that the acceleration of growth, which is characteristic from several growth models, is impaired by the virus.

A similar approach was followed to investigate developmental differences between groups. Mean developmental rates were calculated between consecutive DPIs (Figure 7H). Mock ‒inoculated plants showed a higher mean developmental rate from 5 DPI to the end of the experiment. The Mean Developmental Acceleration (Figure 7I) showed a more complex pattern: Mock-inoculated plants peaked at 5 DPI and after that, a deceleration of development was detected until 12 DPI. In infected plants, though, acceleration reached a maximum at 6 DPI and then sharply decreased towards more negative values than control plants, indicating a relative stagnation in morphological change. At 12 DPI ORMV induced a less negative value of Mean Acceleration of development than controls, although its velocity remained lower ((Figure 7H-I).

Taken together, these results show that ORMV impacts both growth and development very early after infection. Whereas a direct measure (CS) detected differences between treatments at 6 DPI, more elaborated parameters (AGR, RGR and Mean Developmental Acceleration) allowed differences to be detected as soon as 4 DPI. Growth and developmental patterns differed between treatments in a dissimilar manner: AGR showed differences in growth velocities at 4 DPI, whilst Mean Developmental Rate was clearly different later on. Acceleration graphics (Figure 7G,I) indicated that ORMV has an early effect in decelerating both growth and development, but the latter was more dramatically affected in comparison with relative growth deceleration whose decrease was more or less stepwise. Mock-inoculated plants peaked developmental acceleration at 5 DPI and growth acceleration the following day. ORMV-infected plants peaked developmental acceleration at 6 DPI but lost the subsequent growth acceleration phase (Figure 7G,I). This and other comparisons indicate that ORMV does not just induce delayed growth or morphological change patterns, but a more radical change in the coordination of both parameters.

### Comparison with TuMV infections

As stated earlier, one goal of applying the GM approach to the study of Arabidopsis is to make phenotypic comparisons in a more objective and repeatable manner. To this end, the same experimental setup was applied to the study of viral infections of *A. thaliana* with TuMV, an ssRNA+ virus unrelated to ORMV (http://viralzone.expasy.org/). The experiment spanned from 4 to 10 DPI since at 12 DPI excessive curling of some leaves induced by TuMV impaired the correct assignment of landmarks (Supplementary file TuMV 1st.morphoj). Individual datasets were created for each DPI and Procrustes Coordinates extracted. A combined dataset was created and PCA carried on. After outliers exclusion, 27 Mock and 14 TuMV-inoculated plants remained. PCA revealed that PC1 accounted for 49.2% of total variance (much less than the ORMV experiment accounted for) and PC1 plus PC2 accounted for 69.3% of total variance. Again, PC1 mostly separates juveniles from adult rosettes and negative values related predominantly to infected plants which retained a more immature phenotype (Figure 8A). It was supported by the associated wireframe graph which depicts a relative shortening of leaves #11 and #12, similarly to ORMV-infected plants (Figure 3C). PC2 was strongly positively related to infected plants and, similarly to the ORMV case (Figure 3D), reflects the widening of the angle between leaves #9 and #10. PCs 3 and 4 (Figure 8B-C) accounted for 17.7% of total variance and are mainly negatively related to TuMV infection. Discriminant Analysis (Figure 8D-E) showed that, similarly as observed with ORMV, group means were statistically significantly different starting from 5 DPI. Wireframe graphs also evidenced a strong relative shortening of the petioles, similarly to that had been found under ORMV infections (Figure 4F,I), indicating that more compact rosettes are a common outcome of these viral infections. Discriminant power was slightly higher for almost all DPIs in the case of TuMV (Supplementary Table 5, Supplementary Table 3). Moreover, Procrustes Distances were higher for every DPI in the case of TuMV, which induced a Procrustes separation at 8 DPI only matched at 12 DPI for ORMV-infected plants (Supplementary Table 5, Table 1). These results suggest that TuMV is a more severe virus than ORMV is in Arabidopsis, since it induces a more pronounced departure from Mock mean shape.

**Figure 8.**
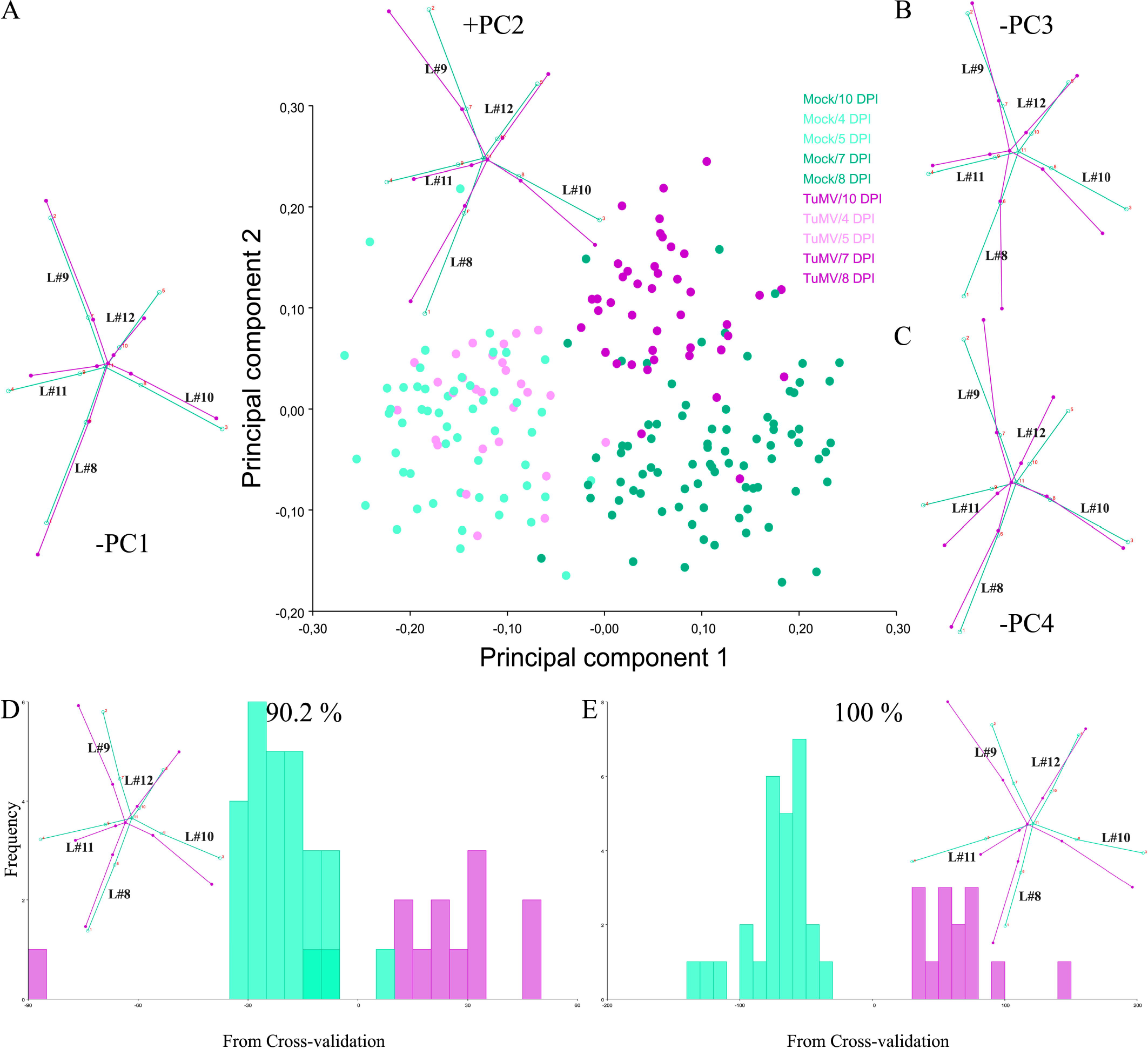
Summary of GM analyses for TuMV-infected plants. (A-C) Shape variation between specimens. (A) PCA scatterplot (PC1 vs. PC2). Pale dots = juvenile (4-5 DPI) plants. Dark dots = mature (7-10 DPI) plants. Wireframe graphs from starting (average) shape (bluish green) to target shape (reddish purple) corresponding to −PC1 (to the left) and +PC2 (top) are included. (B-C) Wireframes for-PC3 and −PC4, respectively. (D-E) Frequencies of jackknifed discriminant scores for 7 and 10 DPI respectively, with wireframes depicting shape changes included. Wireframes show starting shape (mock = bluish green) to the target shape (TuMV = reddish purple). Shape change is magnified 2X.

PTA supported this evidence: A subset of 4-10 DPI datasets were selected to compare ORMV with TuMV infections (Table 2, Figure 9A-B). Whilst trajectory size difference (*MD*_Mock,TuMV_) was similar to the obtained with ORMV, the multivariate angle (*θ*_Mock,TuMV_) that separates infected form healthy trajectories more than doubled that of ORMV. Shape differences (D_*Shape*Mock, TuMV_) between trajectories almost doubled. The majority of the other measures indicated a slower rate of shape change relative to Mock plants, similarly to ORMV infection, but relatively less marked (Table 2). To visualize and compare shape changes, transformation grids with Jacobian expansion factors and lollipops were done in PAST for 10 DPI plants (Figure 9C-D). Both viruses induced relative contraction of the rosette around leaf #11 (the most affected), but TuMV induced more severe deformations. To confirm these results and to test for the reproducibility of the analysis, an independent experiment of TuMV infection was executed (Supplementary file TuMV 2nd.morphoj). PTA analyses were run and trajectory attributes compared (Table 2). There were obtained very similar results relative to the first TuMV experiment.

**Figure 9.**
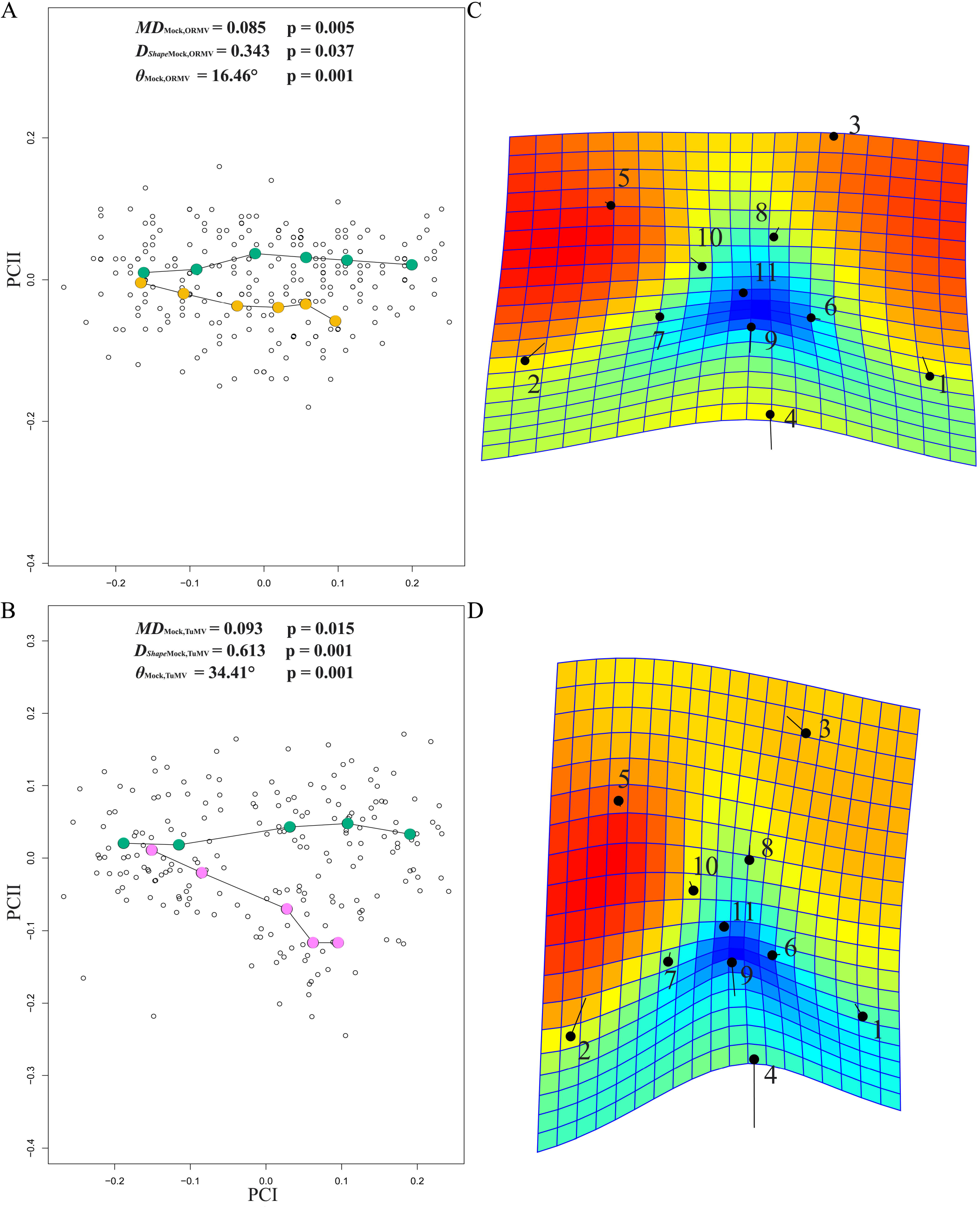
Comparison of virus severity. PC plots of PTA for (A) ORMV- and (B) TuMV-infected plants, compared with Mock-inoculated plants (4-10 DPI). PTA parameters are shown (see main text). Transformation grids with lollipops and Jacobian expansion factors were executed in PAST for (C) ORMV- and (D) TuMV-infected plants depicting shape change from controls at 10 DPI. Jacobian expansion factors indicate expansions of the grid (yellow to orange red for factors > 1) or contractions (blue for factors between 0 and 1). Lollipops indicate target position of landmarks with dots. Leaf #11 (landmarks 4 and 9) is positioned at the bottom.

Together, these results indicated that both TuMV and ORMV induced relative developmental arrest as well as shape change, but symptoms triggered by ORMV are mainly driven by developmental arrest whereas TuMV also promotes shape change in a relatively higher extent, thus impacting more strongly on overall shape.

## Discussion

Here, several standard GM tools were applied to the study and comparison of morphological changes induced in Arabidopsis by viral infections. GM analysis is a powerful approach due both to its statistical toolbox and its appealing visual analysis of shape change. By conceptually separating size and shape, making them mathematically orthogonal, both factors that determine form could be separately analyzed. Thus, the effect of ORMV infection was detected earlier on shape and the derived measures of size (Tables 1-3, Figure 7E,F) than in size itself (Figure 7D). GM analysis greatly outperformed diagnosis when compared against expert human eye (Supplementary Table 3). The effect of time on shape was more pronounced than that of treatment, since the former was detected earlier (Tables 2 and 5). This was particularly the case for control rosettes, reflecting that normal rosette development is not a scaling up of previous shapes but a relative displacement of newly developed structures, a process that is somewhat impaired by ORMV, which induced the retention of a more juvenile-like phenotype (Figure 3).

Normal allometric growth comprised a lengthening of petioles and laminae of new leaves (#11 and 12) relative to older ones (Figure 5H-I). This process was reversed by ORMV, which also distorted the normal angle of approximately 137.5° between successive leaves. As a result, leaves #9 and 10 bended towards leaves #8 and 11, which in turn came close together, bending towards the inoculated leaf (#3) that is situated middle way between them (Figure 4F,I). TuMV provoked similar outcomes (Figure 8) but the effect seemed stronger, not only regarding the distorted inter-leaves angle, but for the relative contraction of leaf #11 respect to all remaining leaves, including #12 (Figure 8E, Figure 9C-D). Taking into account the source-to-sink nature of viral movement by phloem (Manacorda et al., 2013) and its radial structure (Taiz and Zeiger, 2006) it could be hypothesized that virus or viral-induced hormones are distributed through the rosette in such a way that they inhibit proximal systemic growth. These kind of data-based hypothesis is one desired outcome of the application of GM tools (Zelditch et al., 2012) in particular and phenotyping in general. Future work should test this hypothesis by means of comparing cell number or size in distal and proximal parts of systemic leaves, or the effect growth hormones and mutants have in these parameters.

Both viruses diminished shape change, constraining virus-infected rosettes to smaller regions of multivariate morphospace (Supplementary Table 1, Table 2 and Figures 6-9). Ontogeny (the development or course of development of an individual organism) is a genetically-based endogenous process but can be altered by the environment (Gould, 1977). Here, the departure of normal ontogenetic development is induced by both viruses. The consequences of this departure should be analyzed by further work measuring relevant traits.

The availability of a standard measurement unit of shape change (Procrustes distance) allowed to compare ORMV- and TuMV- induced shape changes relative to the departure from healthy control shapes (Figure 9A-B, Tables 2-4 and 6) and objectively rank symptoms severity. Besides, visualization tools aided to identify were the shape change differences allocated in the rosette (Figure 4C, F, I, Figure 8D-E and Figure 9C-D). In sum, it was concluded that TuMV impacts more strongly on Arabidopsis rosette shape than ORMV does. Trajectory and density parameters could be also used to compare developmental phenotypic plasticity (a term generally used to summarize how a given group responds to a series of different environmental conditions by producing an array of phenotypes (Pigliucci, 2004)). Multivariate reaction norms could be then obtained, using shape variables but also controlling for other variables (size, external factors) and weighting their interaction. This would enrich the description of phenotypes whilst offering a solid basis for comparisons.

As superior organisms, plants have complex shapes that experience complex changes throughout their life spans, particularly when exposed to severe stresses that modify the route of ongoing development. Regarding so, their complex phenotypes are difficult to encompass in all their extent by using only one technique, regardless of its descriptive or statistical power. This note is of importance when evaluating the capabilities and limitations of the GM tools presented here. For example, whereas we showed that ORMV significantly impacts rosette shape from 5 DPI and beyond (Supplementary Table 2 and Table 1), and learnt from the wireframes (Fig. 4C, F, I) that some laminae and (mostly) petioles become relatively shorter under ORMV infection, no particular statistical statement could be done about these discrete phenotypic outcomes. Rather, if these questions were to be specifically addressed, other measures (such as direct measures of petioles´ length) should have been taken. GM analyses performed here pointed to overall shape (and size) changes, and visualization tools could serve as guides to further study the putative underlying mechanisms involved if required. Also, as pointed out above, landmarks analyses come with the limitation of not being capable of extrapolate results to the regions between them without uncertainty. Because of that, the selection of a specific set of landmarks (covering the region of interest) must be well stated at the beginning of the experiment and be sound to study the problem of interest. As with any other technique, caution is needed when interpreting the results in order to consider its limitations.

After the genomic revolution, there is a need of objective, reproducible, and accurate assessments of morphology as a critical missing link to supporting phenomics (Punyasena and Smith, 2014). The use of GM allows standardizing deviations from controls in a consistent, objective manner. At the core of these conceptual framework is the GPA, which permits to compare shapes in Procrustes units of distance.

The examples given in this work are necessarily limited, but other applications could be easily envisioned: as the choice of landmarks placement is arbitrary on a given structure, other experimental setups could place them differently to study different stages of growth or other anatomical regions of interest. Importantly, this technique is not a competitor but a possible complementation of newly developed automated platforms for rosette segmentation. It is now possible for some platforms to identify the tip of leaves, the center of the rosette and the intersection between lamina and petiole (Apelt et al., 2015; Tessmer et al., 2013), thus giving the landmarks used in this study and its coordinates, automatically.

Moreover, the same software used in this work permits GM 3D image analysis, therefore allowing the study of plant species with a more complex architecture.

100 years after the revolutionary vision of D´Arcy Thompson´s transformation grids and more than 40 years since the beginning of the revolution in morphometrics, GM application for plant phenotyping is starting to develop (Gómez et al., 2016; Viscosi, 2015; Viscosi and Cardini, 2011) and the plant model species Arabidopsis thaliana should benefit from it.

## ACKNOWLEDGEMENTS

We thanks Dr Flora Sánchez and Dr Fernando Ponz for the kind gift of ORMV and TuMV virus. We thank Dr. Ken Kobayashi and Nicolás Carlotto for human treatment assignments of plants in the Discriminant Analysis comparison, Dra. Valeria Carreira for critical reading of the manuscript and Mariano Manacorda for assistance in adapting figure colours to the Color Universal Design for accessibility to colour-blind people. This research was supported by PICT 2014-1163 from Agencia Nacional de Promotión Científica y Tecnológica (ANPCyT) and by project PE 1131022 (INTA). The authors declare that they have no conflict of interests.

## Appendix S1. Description of the Durbin-Watson panel test

Supplementary Figure 1. Graphical assessment of the Tangent shape space approximation. Scatterplots of distances in the tangent space against Procrustes distances (geodesic distances in radians) for (A) Mock-inoculated plants, (B) ORMV-infected plants and (C) all plants, over all DPIs. A blue line is plotted to show a slope of 1 through the origin. Then a least-squares regression line through the origin is shown in red (for data in which the variation in shape is small this will hide the blue line).

Supplementary Figure 2. Wireframes depicting shape change associated with −PC1 values from 3 to 12 DPI (A-H). Green = starting (average) shape; red = target shape. No magnification was applied.

Supplementary Table 1. Summary statistics for the comparisons between Tangent (Euclidean) and Procrustes shape distances from average shapes and for regression slopes and correlations between the two distances.

Supplementary Table 2. Summary of centroid size and shape variation. Hierarchical sum of squares ANOVA. Main effect: Treatments; random factors: Individuals (ID), Digitization. SS, MS and df refer respectively to sum of squares, mean sum of squares (i.e., SS divided by df) and degrees of freedom. Error1 = Measurement error.

Supplementary Table 3. Classification/misclassification tables from DA for each DPI and human observers for 7 DPI.

Supplementary Table 4. Statistical comparisons of intra-treatment shape changes across the ORMV experiment. Holm´s-Bonferroni sequential correction at α=0.05.

Supplementary Table 5. Discriminant Analysis for TuMV. Statistical tests for differences between means of treatments at each DPI from DA (with permutation tests with 1000 random runs) and classification/misclassification tables for each DPI.

